# ERF transcription factor regulons underpin growth-defence trade-off under acute heat stress in rice seedlings

**DOI:** 10.1101/2025.04.21.649784

**Authors:** Akshay U Nair, Shubham Vishwakarma, Titir Guha, Rajasekhar Varma Kadumuri, Felix B. Fritschi, Sreenivas Chavali, Annapurna Devi Allu

## Abstract

- Rice, a staple cereal crop, faces significant threats from rising temperatures, affecting all growth stages including early seedling establishment. Despite being critical in determining overall growth and productivity, response to heat stress during the early seedling stage remains understudied. This research aimed to assess the impact of acute heat stress on rice seedlings and unravel underlying molecular mechanisms.
- Rice seedlings were exposed to varying intensities and durations of heat stress to determine a critical threshold affecting growth. To elucidate the transcription factor (TF)-mediated regulatory mechanisms and their functional interactome in response to stress, transcriptomic analysis of shoots and roots exposed to acute heat stress was performed.
- Transcriptome analysis unveiled a comprehensive TF-target regulatory map for shoots and roots, potentially involved in the modulation of growth-defence trade-off in response to acute heat stress. Ethylene Responsive Factors (ERFs) emerged as central regulators, with phytohormones ethylene and jasmonic acid acting as upstream modulators. Pre-treatment with these phytohormones alleviated the adverse effects of heat stress.
- This study uncovers key molecular mechanisms governing rice seedling responses to acute heat stress involving ERFs-hormonal interactions. Modulating these core regulators presents a promising strategy to enhance heat resilience, addressing global food security amid rising temperatures.

## Introduction

The climate crisis is intensifying, posing challenges to all forms of life (Calvin *et al*., 2023). Plants must endure adverse conditions that often affect their growth, productivity, and even survival (Bakery *et al*., 2024). Over the last century, the global average air temperature has increased by 0.5°C and is projected to rise by an additional 1.5°C – 4.5°C by 2100, which will increase the prevalence of heat stress and significantly threaten crop productivity in many parts of the world (Calvin *et al*., 2023; Bakery *et al*., 2024). Rice (*Oryza sativa* L.), a staple food crop for more than one-third of the world’s population, is highly sensitive to temperature fluctuations (Wu *et al*., 2023). It thrives at a day-night temperature range of 28°C – 22°C (Bakery *et al*., 2024), and temperatures above 35°C disrupt its growth and development, particularly during the reproductive phase (Begcy *et al*., 2018). India contributes approximately 26% of global rice production, underscoring its crucial role in ensuring global food security. However, tropical countries like India are frequently prone to detrimental high-temperature events (Ravindra *et al*., 2024). Notably, rising maximum daily temperatures caused by short-term heat waves, particularly during the onset of the Kharif season, the major rice cultivation season in India, pose a significant threat to rice cultivation (**Fig. S1a; Table S1.1**).

Plants possess mechanisms to assess ambient temperature and modulate their physiological processes (Wu *et al*., 2023). In response to high temperature, they mount heat shock response (HSR) by activating several genes, such as heat shock proteins (HSPs), ROS scavenging enzymes, and genes involved in hormone and metabolic pathways. Timely activation of HSR is crucial for plant survival and post-stress recovery (Wu *et al*., 2023; Bakery *et al*., 2024). Transcription factors (TFs), as major regulators of stress-induced gene expression changes (Hoang *et al*., 2017), play a key role in HSRs (Guo *et al*., 2016). Multiple TFs and their downstream targets have been identified to influence stress response outcomes (Ohama *et al*., 2017; Chao *et al*., 2017; Albertos *et al*., 2022; Alshareef *et al*., 2022; Tan *et al*., 2023). But, more often, these TF regulations are portrayed as an isolated process, where a TF acts alone or in combination with a few other TFs to control downstream gene expression. Hence, investigating the transcriptional programs regulated by stress responsive-TFs is critical to gaining insights into the molecular mechanisms underlying plant growth and survival under stress (Wang *et al*., 2023; Li *et al*., 2023). Previous studies on rice heat stress majorly focused on the reproductive stage due to its critical role in determining grain yield (Jagadish *et al*., 2010; Hasanuzzaman *et al*., 2013). However, early seedling developmental stages are vulnerable to heat stress, which can significantly impact overall plant growth and productivity (Xiang & Rathinasabapathi, 2022). Despite this, very little is known about seedling responses to heat stress (Begcy *et al*., 2018; Aryan *et al*., 2022), which is a frequent challenge during the rice cultivation seasons in tropical/subtropical regions (Wassmann *et al*., 2009; Huang *et al*., 2017).

To address this, here we first investigated the effects of varying temperature regimes on the growth and survival of rice during the early seedling developmental stage and identified temperatures that inhibit seedling growth. Using transcriptome analysis and reconstructing gene-regulatory networks, we identified key transcriptional programs that drive the response of rice seedlings to short-term, growth-inhibiting heat stress. Such extensive, integrative analyses unveiled that the ethylene/jasmonic acid-mediated regulation of Ethylene Response Factors (*ERFs*) play a pivotal role in modulating the growth-defence trade-off under heat stress during seedling development in rice. Furthermore, we show that targeting upstream mechanisms (ethylene/jasmonic acid response) that modulate the regulatory *ERFs* could serve as a promising approach to optimise developmental decisions and promote heat stress tolerance in rice.

## Results

### Short-term acute heat stress perturbs cellular redox homeostasis, hindering seedling growth

To examine the impact of short-term high temperatures on the growth and survival of rice seedlings, we exposed five-day-old seedlings to elevated temperatures of 35, 40, 45, or 50°C for 2 or 4 hours (h), while the control seedlings were grown at an ambient temperature (28°C). Exposure to temperatures up to 40°C for 2 and 4 h did not hinder seedling growth (**Fig. 1a**). In fact, the seedlings exposed to 35°C for 4 h or 40°C for 2 h displayed enhanced shoot growth (**Fig. 1b**), a phenomenon referred to as thermomorphogenesis (Casal & Balasubramanian, 2019). However, exposure to 45°C for two hours significantly reduced seedling growth, which was more pronounced with increased duration (4 h) (**Fig. 1a,b**). A significant reduction in root growth was also observed in seedlings exposed to 45°C for 4 h (**Fig. 1c**). Notably, a further 5°C increase in temperature (50°C) was lethal to the seedlings (data not shown). These results indicate that 45°C for 4 h is a short-term heat stress regime (hereafter referred to as heat stress) that significantly impedes seedling growth.

**Figure 1.**
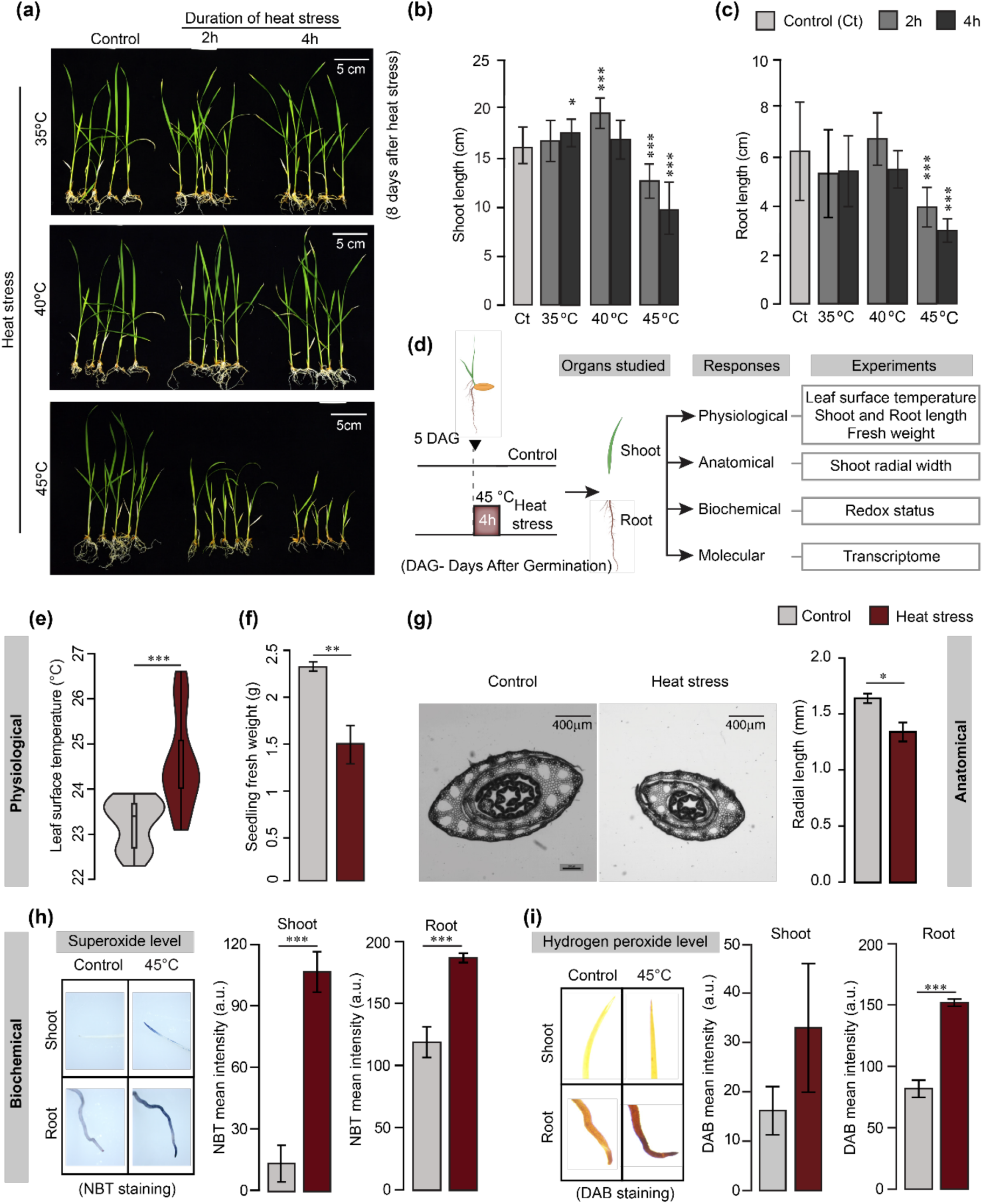
Short-term acute heat stress significantly impairs growth in rice seedlings. **(a)** Rice seedlings (five days after germination) exposed to 35°C, 40°C, or 45°C for 2 or 4 h. Untreated seedlings grown at optimal conditions (28°C) served as control. Scale bars indicate 5 cm. Bars represent mean shoot **(b)** and root **(c)** length (*n* = 15 seedlings) for control and heat-stressed plants (2 and 4 h after exposure at each temperature). Measurements were recorded eight days post-treatment. Error bars represent standard deviation. **(d)** Schema representing an overview of the experimental design. Five-day-old seedlings were exposed to either control (28°C) or heat stress (45°C). Post heat stress, seedlings were sampled at different time points for physiological, anatomical, biochemical, and molecular analyses. **(e)** Leaf surface temperature under control (C) and immediately post heat stress (45°C) conditions (*n* = 18 seedlings; median values along with interquartile ranges are represented). **(f)** Whole seedling fresh weight in response to control and heat stress conditions measured eight days post-heat stress (*n* = 15 seedlings). **(g)** Transverse cross-section images of shoots from control and heat-stressed plants show differences in the radial width. Scale bars = 400 µm (*n* = 8 seedlings). **(h)** Nitroblue tetrazolium chloride (NBT) staining for superoxide in shoot and root. **(i)** Di-amino benzidine (DAB) staining for hydrogen peroxide (H₂O₂) accumulation in shoot and root. The stain colour intensity is given in arbitrary units (a.u.). (*n* = 8 seedlings; mean ± SD is shown). Statistical analyses were performed for all the treatments by comparing them with the control, using Student’s t-test. *P < 0.05; **P < 0.005; ***P < 0.0005.

To understand the mechanisms underlying the seedling response to heat stress, we examined the plant responses at different levels of biological complexity, namely physiological, anatomical, biochemical and molecular (**Fig. 1d**). As plants modulate their energy flux and temperature by dissipating heat energy and increasing evaporative cooling (Gauthey *et al*., 2024), we first checked the leaf surface temperature immediately post-heat stress. Seedlings under heat stress displayed significantly higher leaf surface temperature compared to the untreated controls (**Fig. 1e**), suggesting perturbed leaf transpirational cooling. The impact of heat stress on the growth correlated with the overall decrease in fresh weight of the seedlings (36% reduction; **Fig. 1f**) and a decrease in shoot radial width (22% reduction; **Fig. 1g**) as measured eight days post-heat stress. Reduced radial width under heat stress may be due to the delayed phyllochrone (**Fig. 1g**).

The observed heat-induced morpho-physiological changes may be driven by altered redox homeostasis (Suzuki & Katano, 2018; Huang *et al*., 2019). Hence, we next quantified reactive oxygen species (ROS: superoxide: O_2_^−^ and hydrogen peroxide: H_2_O_2_) levels and their counteractive antioxidant enzymes (superoxide dismutase: SOD and guaiacol peroxidase: GPOX) at different time intervals (0h, 12h and 24h) post-heat stress. Upon heat stress, seedlings displayed elevated O_2_^−^ and H_2_O_2_ content in both shoot and root (**Fig. 1h,i; S1b**). In the shoot, although H_2_O_2_ content was increased immediately post-heat stress (0 h), SOD activity was not impacted at that timepoint, but was significantly greater than in the control at 12 h and 24 h post-heat stress. (**Fig. S1c**). In contrast, SOD activity in roots was not influenced by the heat treatment at any of the three time points (**Fig. S1d**). In contrast, the GPOX activity was higher in both organs immediately post-stress compared to the control but not after 12 h and 24 h (**Fig. S1c,d**). As heat stress and the associated ROS burst often lead to oxidative stress, we also checked the levels of proline, an osmoprotectant with often increased levels under heat stress (Iqbal *et al*., 2019). We observed a heat-induced initial spike in proline content in both organs (0 h), but no difference in the shoot and significantly reduced concentrations in the root at later time points (**Fig. S1c,d**). Although proline is known to maintain osmotic balance in cellular organelles, high proline content for prolonged periods can be lethal (Iqbal *et al*., 2019), which likely explains why seedlings exhibited reduced root proline content at later time points post-heat stress. These results indicate that acute heat stress perturbs various cellular processes, impairing seedling growth. Presumably, the mounted response mechanisms aided in the seedling survival but were not efficient enough to completely mitigate the impact of heat stress on growth.

### Transcriptome analysis reveals growth defence trade-off in both shoot and root

We analysed the transcriptomes of the shoot and root to identify the regulatory mechanisms that modulate the acute heat stress-induced response. Interestingly, among the overall differentially expressed genes (DEGs), more genes were upregulated than downregulated in both shoot (∼6% more) and root (∼9% more) (**Fig. 2a; Table S1.2**). While the majority of the common DEGs (those overlapping between shoot and root) were upregulated genes, the DEGs unique to shoot or root comprised more downregulated genes (**Fig. 2b**). The overlap of DEGs (2636 genes) between shoot and root was significant (Z-score =75.34; **Fig. 2c**). Next, we checked the contribution of these common and unique sets of genes to acute heat stress responses. For this, we further categorised the DEGs into different sets (up-/down-regulated in shoot/root) and performed gene ontology (GO) enrichment analysis (**Fig. 2d, S2**). For both organs, we observed significantly enriched biological processes, such as response to heat, regulation of RNA biosynthetic process, transcription for the upregulated genes, and, ion transport, chemical homeostasis and cell wall organization for the downregulated genes. Biological processes such as response to oxygen-containing compound/hydrogen peroxide, protein folding, carbohydrate metabolic process, and developmental process were significant in the shoot. This indicates activated ROS detoxification mechanisms and aligns with the relatively lower hydrogen peroxide levels observed in the shoot than in the root (**Fig. 1i, S1b-d**). Root-specific processes represented downregulated genes, that included the response to auxin, transmembrane transport and lignin-/phenypropanoid-catabolic process. While auxin homeostasis is enriched for downregulated genes in the roots, shoots displayed regulation of the jasmonic acid (JA) signalling pathway for the upregulated genes. While, in both shoot and root, the upregulated genes were enriched in stress response-related processes, the downregulated genes were associated with several growth-related processes (**Fig. 2d, S2**). These observations support a potential stress response-growth trade-off upon acute heat stress, with these genes representing the underlying molecular players.

**Figure 2.**
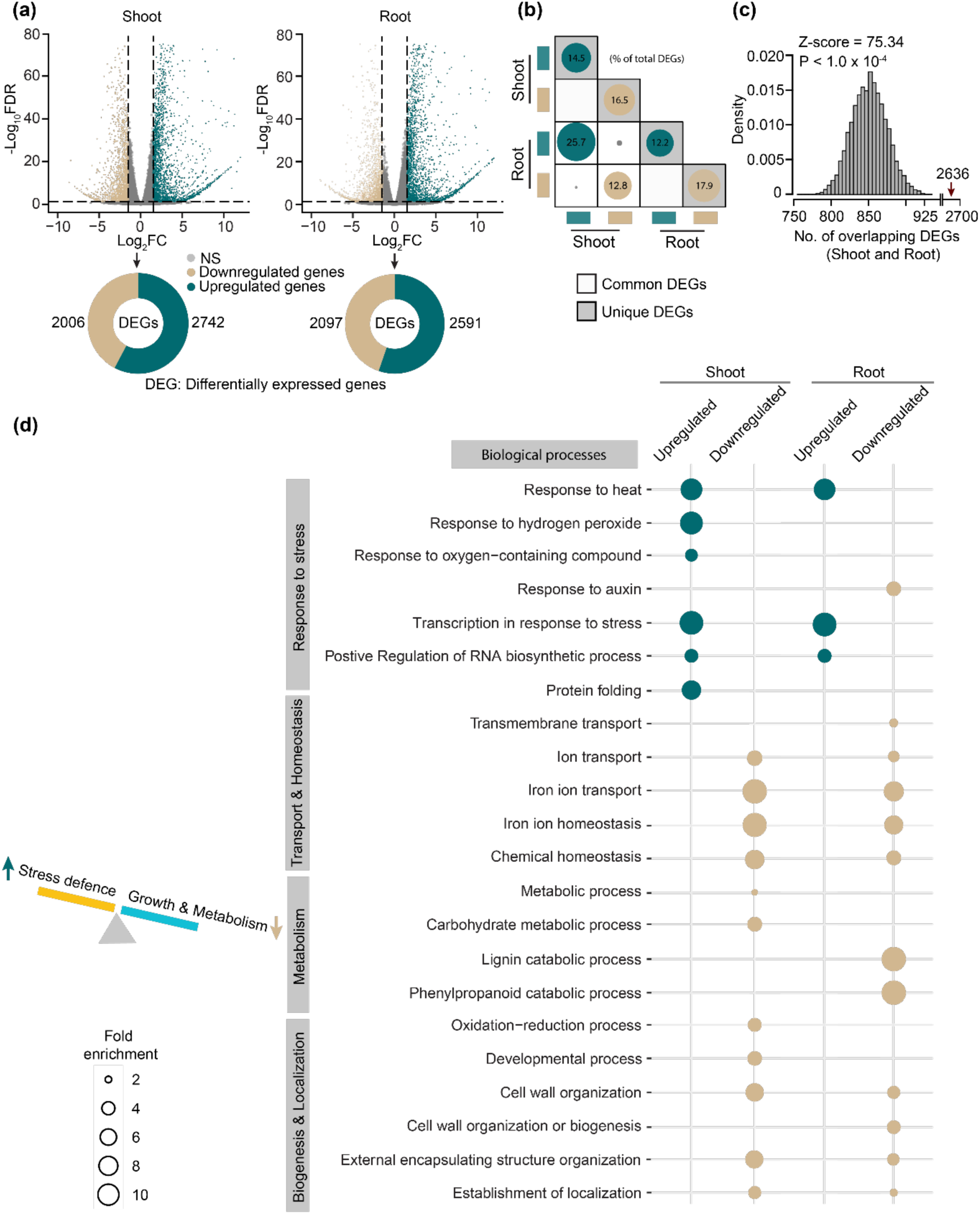
Conserved and distinct transcriptional responses across shoot and root under heat stress. **(a)** Volcano plots showing differentially expressed genes (DEGs) in the shoot and root in response to heat stress. DEGs were identified with thresholds of log_2_ fold change ≥ 1.5 or ≤ – 1.5 and false discovery rate (FDR) < 0.05. Upregulated genes are shown in green and downregulated genes are in brown. The pie charts below each Volcano plot indicate the total number of DEGs for the shoot and root. **(b)** Matrix plot displaying the number of DEGs that are common (plain boxes) and unique (grey boxes) to shoot and root. Bubble size indicates the fraction of DEGs under each category. **(c)** Enrichment of overlapping DEGs among root and shoot assessed using permutation testing. The grey histogram represents the random expectation and the red arrow represents the actual observation. **(d)** Enriched biological processes associated with DEGs in the shoot and root. Bubble plot showing up- and down-regulated biological processes. Bubble size corresponds to fold enrichment.

### Diverse transcription factors co-ordinately regulate seedling responses to acute heat stress

In plants, stress-induced gene expression changes are majorly governed by transcription factors (TFs) (Wang *et al*., 2023a; Li *et al*., 2023b). Indeed, we observed that the gene ontology (GO) term ‘transcription in response to stress’ was enriched for upregulated genes in both shoot and root (**Fig. 2d**). A total of 44 out of 56 TF families in rice (according to PlantTFDBv5.0; https://planttfdb.gao-lab.org/) were found to be differentially expressed (DETF) in both organs (**Fig. 3a**). Notably, both shoot and root display substantial diversity in the engagement of TFs in response to heat stress (**Fig. 3a, S3**). The TF families such as *NAC, WRKY, MYB, HSF, bZIP, ERF, bHLH* and *C2H2* were prominent in both shoot and root. Identification of the TFs from families such as *NAC, WRKY, MYB, HSF, bZIP*, and *ERF,* to be involved in acute heat stress response in this study corroborates with previous observations, serving as a validation for our results (Surabhi & Badajena, 2020) (**Fig. 3a**). These families displayed more upregulated TFs in both shoot and root, whereas the *bHLH* and *C2H2* families, which were also prominent TF families, displayed more downregulated TFs in both shoot and root (**Fig. 3a**). To elucidate the TF-mediated regulation of heat stress responses, we reconstructed DETF-target regulatory networks (**Fig. S4, 3b**). For this, we surveyed the promoter regions of the DEGs for DETF binding site enrichment, which allowed us to identify the DETF-target pairs. We estimated the Pearson correlation coefficient and applied a stringent FDR cut-off <0.05, to identify DEG targets that could be potentially regulated by DETFs (**Fig. S4**). The huge number of TF-target pairs (6223 in shoot and 5211 in root) identified (**Table S1.3**) suggests substantial underlying regulatory complexity.

**Figure 3.**
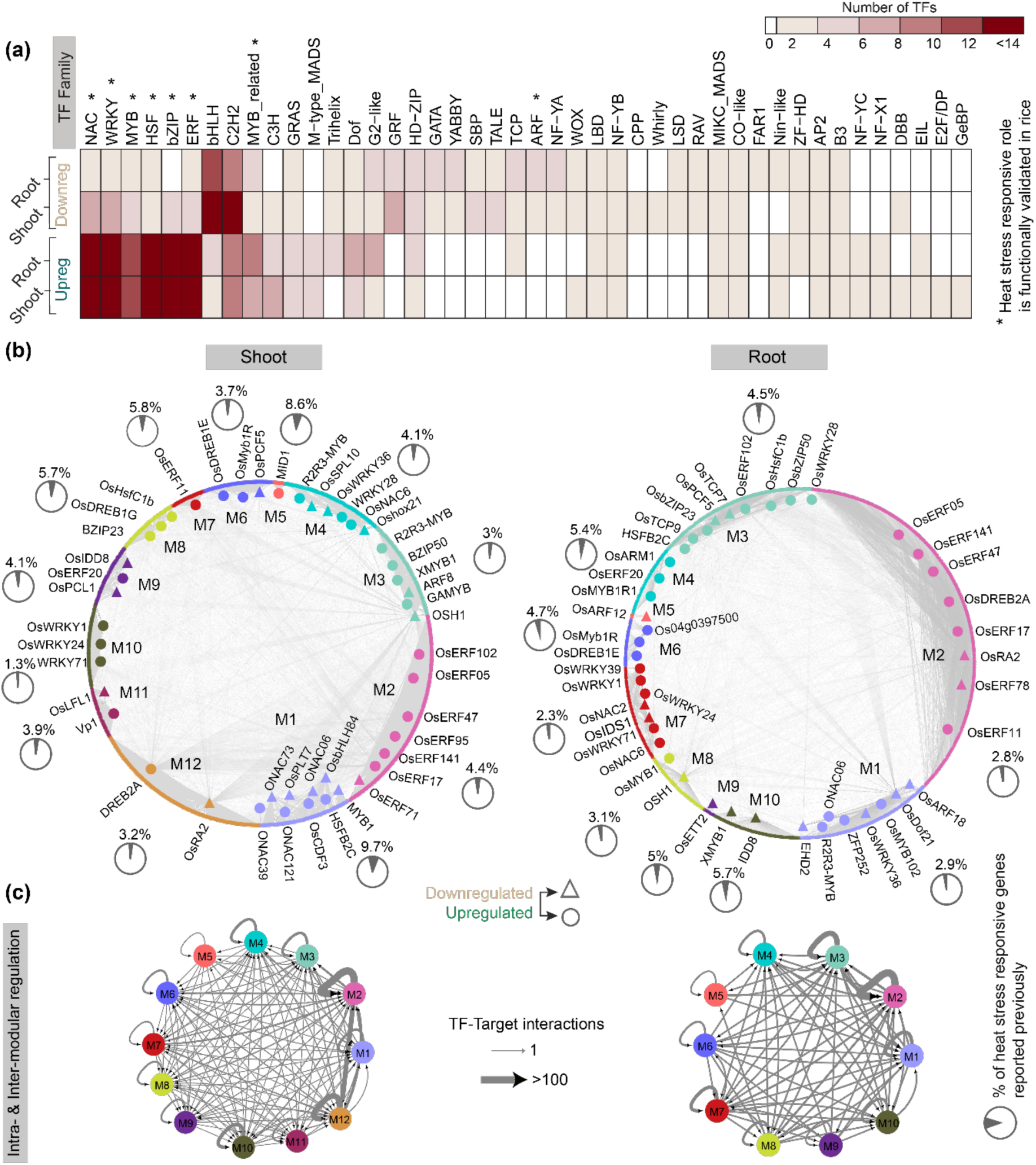
Diverse transcription factors co-ordinately regulate short-term heat stress responses. **(a)** Heatmap showing the number of differentially expressed transcription factors (DETFs) across 44 TF families in shoot and root. Colour intensity represents the number of TFs. **(b)** Gene regulatory networks showing multiple transcription factor (TF)-target modules in both shoot (M1–M12) and root (M1–M10). Each module is colour-coded and highlights key regulatory TFs in them. The percentage of genes previously reported (according to STIFDB2) to be heat stress-responsive is indicated in pie charts for each module (Root module M5 did not have any overlap). Nodes represent genes (circles for up-regulated and triangles for down-regulated, respectively). **(c)** Inter- (between modules) and Intra- (within modules) modular regulations for the entire TF-target network. The density of the edges is proportional to the TF-target interactions.

Transcription factors and their downstream target genes constitute regulatory units, termed ‘regulons,’ which orchestrate various cellular processes and functions (Müller-Dott *et al*., 2023). The extensive number of DETF–target interactions suggests potential crosstalk between DETFs and their regulons. To investigate the regulatory architecture of such crosstalk underlying the acute heat stress response in rice seedlings, we assessed (i) the type of regulons, and (ii) the degree of connectivity within (intra) and between the regulons (inter) by performing topological clustering for DETF-targets of both organs (**Fig. S4, 3b**). We found only a few simple regulons, which represent sets of genes under the control of a single TF (**Table S1.3**), while the majority of the DEGs were part of complex regulons in both shoot (12 modules) and root (10 modules) (**Fig. 3b, S5a**) which included several TFs in each module. Importantly, certain modules of shoot and root had a high representation of a few TF families (at least 50% representation of TFs from a single family within the module). For instance, *ERF* (shoot M2, M7, and M12 modules, root M2 module) and *WRKY* (shoot M10 module, root M7 module) TFs were predominant within several modules. Notably, only a small fraction of genes from the identified TF modules were previously reported to be heat stress-responsive genes (Naika *et al*., 2013). The overlap with previously reported DEGs in heat stress response ranged between 1.3 (module M10) and 9.7% (module M1) in the shoot. In contrast, the overlap in root ranged between 0 (module M5) and 5.7% (module M10) (**Fig. 3b**).

Furthermore, GO enrichment analysis revealed that distinct modules regulate distinct processes, while certain modules can regulate common processes. For instance, modules M6 and M9 of the shoot and M3 of the root potentially regulate stress response, while modules M6, M8, M9 and M10 of the root regulate growth and metabolism. On the contrary M1-M4, M8, M10-M12 of the shoot and M1, M2, M4, M7 of the root regulate both processes (**Fig. S5b**). Most of the enriched biological processes were associated with multiple modules, indicating potential cross-module regulation. Hence, we examined whether these modules are regulated by intra-modular TFs (those from within the module) and/or inter-modular TFs (those from across modules) (**Fig. 3c**). The modules M2 and M12 of the shoot, and M2 and M3 of the root exhibited the highest intramodular regulation (**Fig. 3c**). These modules also demonstrated higher intermodular regulation between them in the respective organs. Strikingly, the shoot M2 and M12 and root M2 modules were found to potentially regulate both growth and stress response (**Fig. S5b**). The distinctive characteristics of these modules, in conjunction with their association with both growth and stress responses, suggest a potential functional role for this subset of transcription factors in driving the heat stress response outcomes.

### ERF regulons are crucial in regulating short-term heat stress responses

To further check the potential functional cohesiveness of these DETF-regulons (modules), we retrieved DEG-functional interactome (using STRINGDB), to identify hub genes for both shoot and root (**Fig. 4; Table S1.4**). In line with the observed DEG GO enrichment (**Fig. 2**), functional interactome also displayed significant enrichment for stress response for the upregulated genes and growth and development for the downregulated genes (**Fig. S6**). Next, we identified the core hub (a cluster of genes having the highest centrality measures) (**Fig. 4a**). The core hub genes are known to potentially play a crucial role in the structural and functional organisation of biological networks (Kim *et al*., 2017; Liu *et al*., 2021; Yin *et al*., 2023a). Genes from shoot modules M2 and M12 and root modules M2 and M3 represented a major proportion of core hub genes among genes from other modules (**Fig. 4b**). Interestingly, we found the ERF family DETFs to be the major regulators of these modules (shoot M2, M12 and root M2; **Fig. 4c**) with a similar pattern of expression in both shoot and root upon heat stress (**Fig. 4d**). Collectively, these findings at the TF-regulon and DEG-functional interactome level, project *ERF* DETFs-*OsERF5, OsERF11*, *OsERF47, OsERF102* and *OsERF141* as potential regulators of heat stress-induced growth and defence responses in rice seedlings.

**Figure 4.**
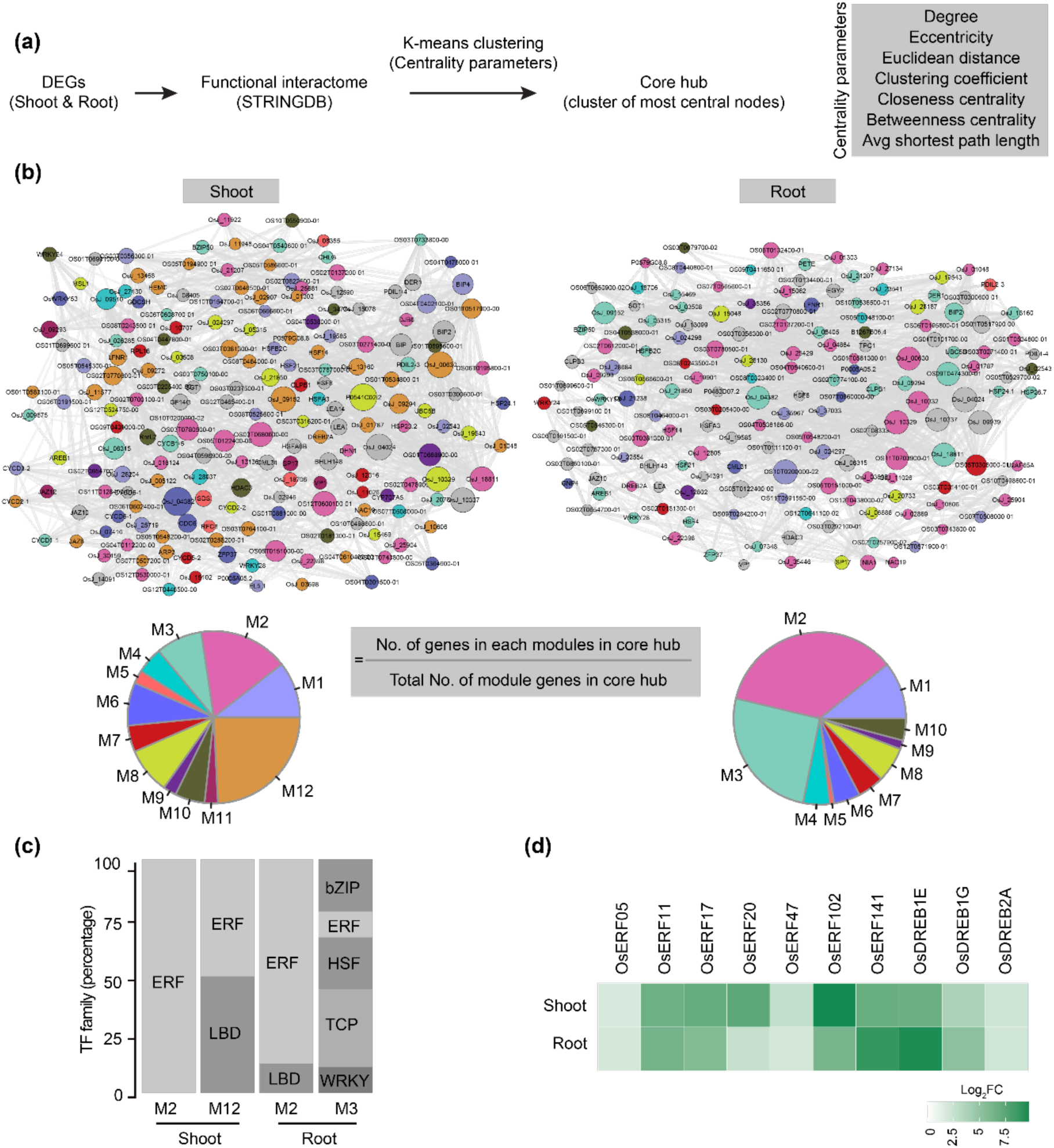
ERF regulons are crucial in regulating short-term heat stress responses. **(a)** Workflow illustrating the identification of core hub of the functional interactome **(b)** Core hub of the shoot and root functional interactome (STRINGDB) network, with various module genes highlighted in respective colours. Pie charts indicate the fraction of genes each TF-target module represents in the total module genes in the core hub. **(c)** Bar plot indicating the percentage of different TF families regulating the M2, M12 shoot modules and M2, M3 root modules. **(d)** The heat map indicates gene expression (Log_2_FC) values of *ERF* TFs in both shoot and root after 4 h of 45°C heat stress compared to control.

### Ethylene- and jasmonic acid pre-treatment modulates the growth-defence trade-off upon heat stress

Given the number of *ERF* DETFs identified as key players, we surmised a regulation that operates upstream to collectively fine-tune their expression, thereby regulating heat stress response outcomes. The phytohormones, ethylene and jasmonic acid (JA) are known to regulate *ERF*s (Müller & Munné-Bosch, 2015). Our data also revealed the upregulation of genes involved in the biosynthesis and signalling of both hormones (**Fig. 5a,b**). Further, measurements of ethylene emission from the rice seedlings exposed to heat stress (45°C for 4h) displayed a significant increase (∼ 3-fold) in ethylene levels compared to the control (**Fig. 5c**). An increase in jasmonic acid level was observed post-heat stress compared to control (**Fig. 5d**). Additionally, we also observed that the ERF-regulated/preponderant modules (M2 in both shoot and root) display enrichment for the ‘ethylene-mediated signalling’ processes (**Fig. S5b)**. These observations prompted us to hypothesize that ethylene and JA may act as key modulators of heat stress response in rice seedlings through the regulation of *ERF*s. To test this, first we assessed the impact of heat stress on seedlings with or without pre-treatment with the ethylene precursor, 1-aminocyclopropane-1-carboxylic acid (ACC) or the JA supplemented as methyl jasmonate (MeJA) (**Fig. 5d**), hereafter referred to as ethylene and JA. As indicated by shoot length and radial width, both pre-treatments significantly reduced the impact of heat stress on seedling growth when compared to seedlings directly exposed to heat stress (**Fig. 5e-i**). We further examined if ethylene-/JA-induced heat stress response is mediated by the subset of regulatory *ERFs*-*OsERF5, OsERF11*, *OsERF47, OsERF102* and *OsERF141* identified here. For this, we profiled the expression of *ERF*s in seedlings pre-treated with ethylene and JA in the presence or absence of heat stress (**Figure 5j**). Ethylene and JA treatment induced the expression of almost all the *ERFs*, except *OsERF5* whose expression was downregulated in response to JA. Interestingly, the expression levels of all tested *ERFs* in the pre-treated seedlings, regardless of whether they were subsequently exposed to heat stress, were largely similar to those observed in plants subjected to heat stress alone. This, together with induced levels of ethylene and JA upon heat stress (**Figure 5c,d**), suggests that heat stress-induced *ERF* expression might be regulated by ethylene/JA signalling. Taken together, these results suggest that heat stress-induced ethylene and JA may act upstream of the regulatory *ERFs*, which when triggered through the pre-treatment of these hormones, enhances heat stress tolerance and limits impact on growth.

**Figure 5.**
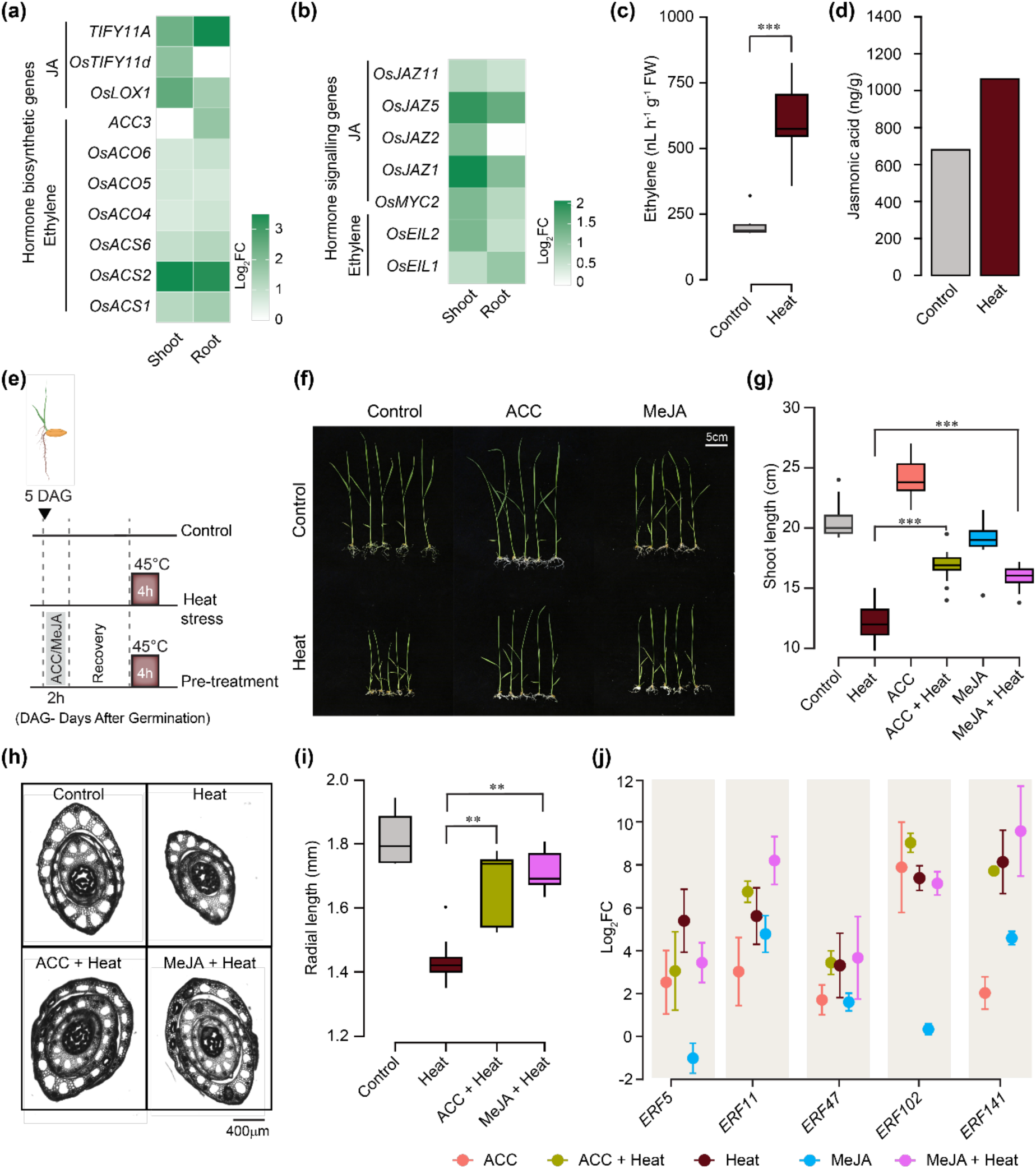
Ethylene- and jasmonic acid pre-treatment modulates the growth-stress trade-off upon heat stress. Heatmap showing expression levels (Log_2_FC) of key ethylene and jasmonic acid **(a)** biosynthesis genes and **(b)** signalling genes in the shoot and root after exposure to 4 h of 45°C. Green indicates upregulated expression, while white indicates minimal or no expression changes. **(c)** Box plot showing the ethylene emission from control and 4 h heat-stressed seedlings. (*n* = 6 biological replicates) **(d)** Bar plot showing the jasmonic acid quantification. (*n* = 2 biological replicates). **(e)** Schema representing treatment regime used to test the effects of ACC (ethylene precursor) or MeJA (jasmonic acid) on rice seedling responses to heat stress. Five days after germination (DAG) seedlings were treated with ACC or MeJA for 2 hours and, following 2 hours of recovery, were exposed to heat stress at 45°C for 4 hours. **(f)** Representative images of seedlings from control and heat stress treatments with and without hormonal pre-treatment. Scale bar = 5 cm. **(g)** Box plot displaying shoot length across control and heat-stressed seedlings, with and without ACC or MeJA pre-treatment (*n*= 15 seedlings). **(h)** Cross-section images of shoots from control, heat-stressed with and without ACC, MeJA pre-treatments. Scale bars = 400 µm. **(i)** Box plot showing shoot radial length; (*n=* 8 seedlings). **(j)** Differential expression of ethylene response factor (*ERF*) genes (*ERF5, ERF11, ERF47, ERF102*, and *ERF141*) across treatments. qRT-PCR data are represented as mean ± SD of three biological replicates, each biological replicate with three technical repeats. Statistical analyses were performed using the Wilcoxon rank sum test. *P < 0.05; **P < 0.005; ***P < 0.0005.

## Discussion

Our planet is experiencing an increasingly warmer and in many locations drier environment, threatening all forms of life (Calvin *et al*., 2023). Alarmingly, between July 2023-24, several parts of the world recorded temperatures above or around 50°C (WMO, 2024; Kacprzyk *et al.,* 2025). The recent scenario of prolonged summers with elevated mean temperatures and sudden heat waves is a significant concern for tropical and subtropical regions, which are among the prime areas of rice cultivation (Vecellio *et al*., 2023; **Fig. S1a**). Despite this, little is known about the heat stress response during early seedling development, which is highly vulnerable to temperature fluctuations as it involves the key transition from self-nourishing to a photosynthetically active state (Facelli, 2008; Han *et al*., 2009; Jung *et al*., 2012; Sarkar *et al*., 2014; Mangrauthia *et al*., 2017). Previous research investigating the impact of heat stress in rice majorly focussed on flowering and grain-filling stages with a duration of exposure from one to more weeks (Yang *et al*., 2015; González-Schain *et al*., 2016; Wang *et al*., 2019). Studies on understanding short-term heat stress response in rice, particularly at the molecular level are limited (Cai *et al*., 2023; He *et al*., 2023). In this study, we documented that short-term acute heat stress adversely affects growth in rice seedlings at the early developmental stage. Exposure to 45°C for 2-4 h impeded seedling growth (hereafter referred to as heat stress; **Fig. 1**). This indicates the vulnerability of rice seedlings to elevated temperatures that closely mimic the recorded temperatures during the initial stages of the rice growing season (WMO, 2024; Kacprzyk *et al*., 2025), particularly in India (**Fig. S1a; Table S1.1**). Upon heat stress, the observed reduction in growth (both in shoot and root) correlated with an increase in leaf surface temperature and higher levels of reactive oxygen species and antioxidant enzymes, reflecting the perturbation and ongoing readjustment of cellular homeostasis under short-term heat stress (**Fig. 1, S1b-d**).

Transcriptome analysis of the shoot and root revealed an activated heat shock response (HSR) with induced expression of heat stress-response genes and transcription factors. Notably, both shoots and roots displayed distinct and common mechanisms with a large overlap in differentially expressed genes (DEGs) and transcription factors (DETFs) (**Fig. 1, 2**). GO enrichment analysis revealed that the several growth-associated biological processes were enriched for downregulated genes, while the stress responses were associated with the upregulated genes. This supports our observation of a growth-defence trade-off in the seedlings. This phenomenon of growth-defence trade-off is well documented in multiple species (Huot *et al*., 2014; He *et al*., 2022), yet the regulatory mechanisms underlying such growth-defence trade-off, particularly in crop plants, remain poorly understood.

Moreover, regulation in biological systems is often complex and may involve multiple regulators (such as TFs, phytohormones, kinases etc.) with intricate crosstalk mechanisms (Yin *et al*., 2023b; Wang *et al*., 2023b). Previous studies involving a muti-omics approach, mapped genome-wide TF-target gene regulatory networks (GRNs), revealing the transcriptional regulation underlying plant responses to various stresses (Wilkins *et al*., 2016; Song *et al*., 2016; Wu *et al*., 2021; Reynoso *et al*., 2022; Sun *et al*., 2022). However, in the context of heat stress in rice, prior research involving transcriptome analysis has predominantly concentrated on the identification of candidate genes or biological processes and pathways (Liu *et al*., 2020; Zhang *et al*., 2022; Yang *et al*., 2022). Our observation of the diversity in DETF families in both shoot and root suggested a complex regulation involved in seedling response to heat stress. To unravel the regulatory complexity, we used a two-tier approach by (i) identifying the TF-target regulons and (ii) overlaying the functional interactome of the DEGs for both organs. This approach revealed an extensive and comprehensive map of regulons in both the shoot and root, with a modest overlap of genes with previous reports that dealt with heat stress response in rice (**Fig. 3b**). This may imply that the seedling response to short but growth-inhibiting heat stress differs from other developmental stages or stress regimes. In line with the observed transcriptional regulation of growth-defence trade-off, the DEGs’ functional interactome network also revealed development and stress responses as down- and upregulated processes, respectively (**Fig. S6**).

Further analysis revealed the regulatory crosstalk within (intra) and across (inter) modules involving multiple DETFs (**Fig. 3**). Interestingly, modules with high intramodular regulation in both shoot (M2 & M12) and root (M2) revealed ERF TFs as key regulators (**Fig. 3**). High intramodular connectivity could indicate a central role for these nodes in a module (Freyre-González *et al*., 2013; Zhou *et al*., 2021; de Silva *et al*., 2022). Furthermore, the overrepresentation of the genes from ERF-regulated modules in the functional interactome core hub of both shoot and root (**Fig. 4**) suggests ERFs as the key regulators of the transcriptional programs that govern the heat stress response in both shoot and root at the early seedling development stage in rice.

In plants, the ERF family of transcription factors are believed to have originated from viruses and bacteria through lateral gene transfer (Licausi *et al*., 2013). ERFs regulate multiple stress responses across different plant species (Dey & Corina Vlot, 2015), and thus are considered versatile regulators. Notably, among three classes of ERFs (AP2/ERF/RAV), the ERFs identified in this study are from the ERF subclass. Lack of validated TF-target datasets (supported by e.g., ChIP-seq, DAP-seq) for rice, including under heat stress, limits the confidence of our TF-regulon predictions, and thus the identification of key TFs. Nevertheless, the ERF subclass TFs were previously reported to regulate heat stress responses in Arabidopsis, which supports our analysis approach in identifying key regulators. For instance, ERF1, ERF95 and ERF97 mediate ethylene pathways and modulate heat stress response by regulating the expression of key *HSPs* and heat shock factors (Huang *et al*., 2023). Stress-induced regulation of *ERFs* involves phytohormones, predominantly ethylene and jasmonic acid (JA). Heat stress is known to induce ethylene and JA in multiple species (Huang *et al*., 2023). Similarly, an elevated level of ethylene and JA was detected immediately post-exposure to acute heat stress in rice seedlings (**Fig. 5**). Ethylene pre-treatment has been shown to enhance thermotolerance in various plant species (Huang *et al*., 2023). However, these studies primarily focused on exploring the phenotypic effects with limited insights into the underlying regulatory mechanisms. As our study identified ERFs as key regulators of the heat stress response, we surmised a core upstream regulatory mechanism involving ethylene/JA governing the ERF-transcriptional programs. Remarkably, pre-treatment with precursor of ethylene/MeJA limited the impact of heat stress on seedling growth, potentially by establishing a primed state mediated through *ERF* DETFs and their associated signalling and transcriptional networks (**Fig. 5**).

In summary, our findings demonstrate that the TF regulons play a crucial role in regulating plant heat stress response outcomes. We present a comprehensive map of heat stress-responsive regulons in the shoot and root of rice seedlings exposed to short-duration, growth-inhibiting heat stress. Studies so far used single-gene/TF-centric methods to tackle growth-defence trade-off and stress resilience. However, modulation of growth under stress involves complex regulation involving multiple TFs (Song *et al*., 2016). This study presents a novel approach to identifying core transcriptional regulators of stress responses, with the potential to boost crop improvement programs. For instance, identifying core TFs and their target genes involved in stress tolerance can lead to the development of molecular markers to efficiently select genotypes with enhanced stress resistance, thereby accelerating breeding programs (Janni *et al*., 2020). The knowledge of these regulatory modules can be directly applied to engineer stress-resilient crops through targeted genetic modification using CRISPR/Cas-mediated precise editing of the core TFs or their binding sites, allowing for the fine-tuning of stress responses (Wang & Niyogi, 2025). Furthermore, we show that targeting the upstream regulatory mechanisms through ‘priming or pre-treatment’ may hold the key to finetuning the growth defence trade-off and offer a sustainable alternative to promote crop resilience.

**Figure 6.**
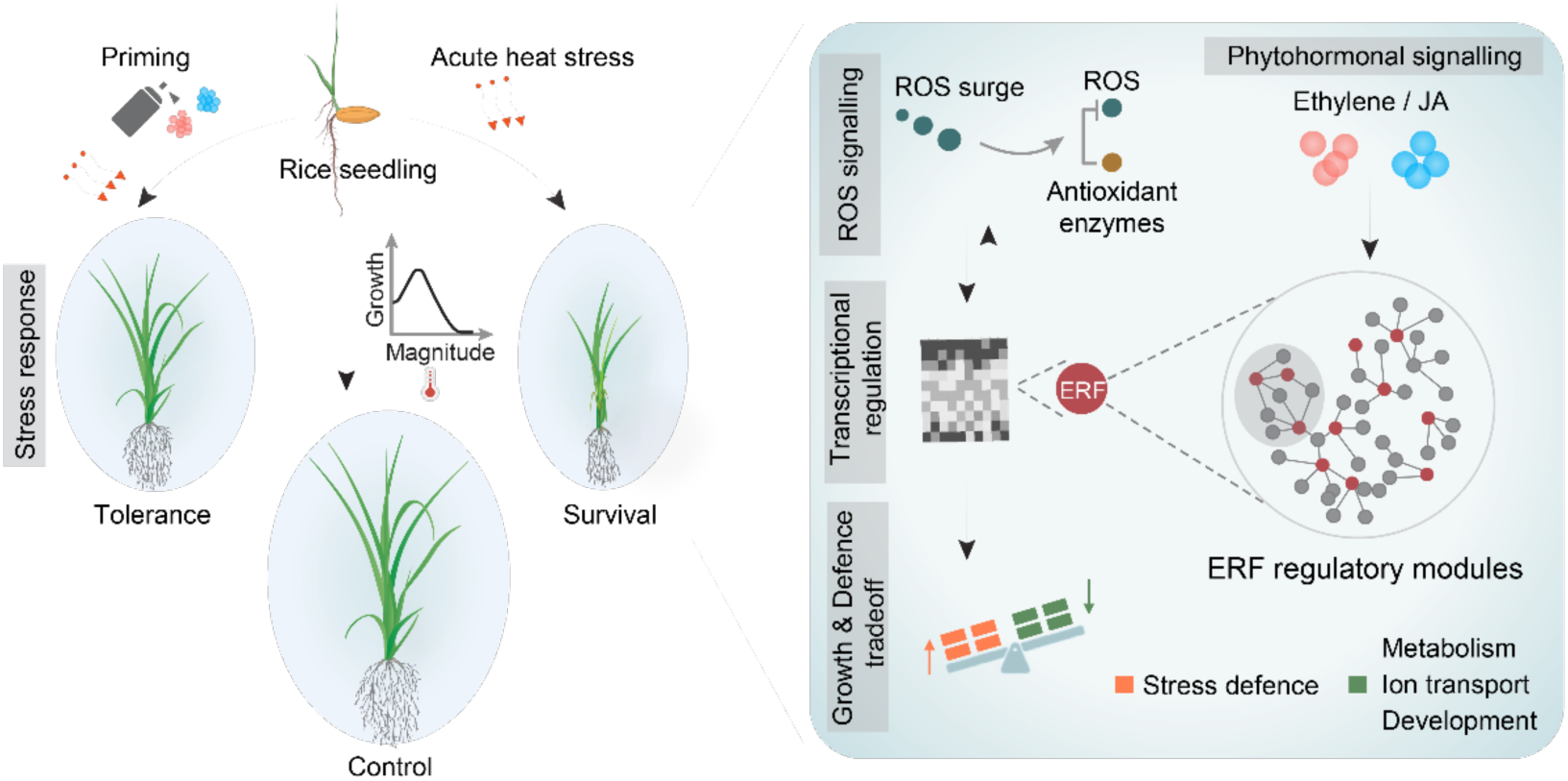
Schema representing molecular and physiological response of rice seedlings to acute heat stress. Rice seedlings, upon exposure to acute heat stress, exhibit a growth penalty accompanied by the activation of stress defence mechanisms. A rapid surge in reactive oxygen species (ROS) triggers antioxidant enzyme activity, while concurrent phytohormonal signalling (ethylene and jasmonic acid) initiates a transcriptional cascade. Central to this cascade is the activation of *Ethylene Responsive Factors* (*ERFs*), which orchestrate downstream regulatory modules involved in modulating the growth-defence trade-off. Notably, pre-treatment with ethylene/JA enhances thermotolerance by regulating *ERF*s. These findings highlight the potential of targeting such transcriptional hubs to confer resilience to heat stress.

## Materials and Methods

### Plant materials and growth conditions

*Oryza sativa* cultivar *Indica v*. N22 seedlings were grown hydroponically in 1X Hoagland solution following a previously described protocol (Hoagland & Arnon, 1950) with slight modifications. Rice seeds were surface-sterilized using 1% sodium hypochlorite for 10 minutes, followed by thorough washing with sterile distilled water. Seedlings (15–20 per jar) were cultivated in phyta-jars with transparent bodies and lids equipped with valves for aeration. The volume of Hoagland solution in each jar was adjusted based on the growth stage of the seedlings, ranging from 40 to 80 mL. The growth conditions were maintained at 14 h light (28°C)/10 h dark (26°C) cycles, with a relative humidity of 65–70%. The light intensity was set at 150–200 μmol m⁻² s⁻¹, and the phyta-jars were kept in a controlled growth room to ensure consistent environmental parameters.

### Treatment conditions

#### Heat stress treatment

Five-day-old seedlings were chosen for heat stress experiments. To optimise the temperature and duration of the heat stress treatment, seedlings were exposed to 35°C, 40°C, and 45°C for 2 and 4 hours (h) (placed in phyta-jars containing 1x Hoagland solution) in a water bath closed with a transparent lid. Air temperature inside the water bath was also monitored using a thermometer, to reach the set water bath temperatures before starting the stress paradigm. Following heat stress, seedlings were allowed to recover under standard growth conditions for 8 days.

### Ethylene and jasmonic acid treatment

For ethylene and jasmonic acid pre-treatment, five-day-old rice seedlings were supplied with 100 μM 1-Aminocyclopropane-1-Carboxylic Acid (ACC; cat-149101) or 10 μM, Methyl jasmonate (MeJA; cat-392707) in Hoagland medium. Pre-treatments were done for 2 h and replaced with Hoagland medium. Post 2 h recovery, seedlings were exposed to heat stress. The other set of seedlings was directly exposed to heat stress without any pre-treatment. Untreated seedlings served as control.

### Physiological measurements

Physiological parameters were measured eight days post-treatment, including shoot length (stem base to the tip of the emerging leaf), root length (length of the primary root) and fresh weight. Immediately post heat stress, leaf surface temperature was assessed using a thermal imaging IR camera (FLIR A325sc), and the data was analysed with FLIR RIRmax software. For shoot anatomical studies, 100 µm-thick transverse sections (2 cm from the base of the shoot) were prepared from control and heat stress seedlings eight days post-stress, using a vibratome (Compresstome® VF-310-0Z) and visualised with a Nikon ECLIPSE Ti2 microscope using a 20X magnification.

### 3,3′-Diaminobenzidine tetrahydrochloride (DAB) staining

To determine the cellular H₂O₂ levels, we used 3,3′-diaminobenzidine tetrahydrochloride (DAB) staining. Measurements were conducted immediately post-heat stress treatment. Intact heat-stressed and control seedlings were vacuum-infiltrated with the staining solution containing 1 mg/mL DAB in 200 mM sodium phosphate buffer (pH 7.5) and 25 µL Tween-20. The samples were incubated in the dark for 24 hours at room temperature (26°C). In turn, chlorophyll was removed by incubating the seedlings in 90% ethanol at 70°C for 30 minutes to 1 hour, or until complete decolorization. Images were captured using a Magnus TZM6 microscope at 2X. Stain intensity was quantified for similar regions of interest of shoot and root from the different treatments using ImageJ software.

### Nitroblue tetrazolium (NBT) staining

Superoxide content was analysed using nitroblue tetrazolium (NBT) staining, immediately post-heat stress. For staining, heat stressed and control intact seedlings were incubated in 1 mg/mL NBT solution prepared in 50 mM sodium phosphate buffer (pH 7.8) for 4 h at room temperature (28°C). After staining, chlorophyll was removed by washing the leaves with a solution of acetic acid:glycerin:ethanol (1:1:4) in a water bath preheated to 85°C, followed by a final wash with glycerin:ethanol (1:4). Images were captured using a Magnus TZM6 microscope at 1X. Stain intensity was quantified for similar regions of interest of shoot and root from the different treatments using ImageJ software.

### Biochemical analysis

To determine the activity of superoxide dismutase (SOD) and guaiacol peroxidase (GPOX) enzymes, shoot and root tissues were sampled at 0, 12, and 24 h post heat treatment regime and immediately frozen in liquid N_2_. Frozen shoot and root tissues (0.1 g FW) were ground using a chilled mortar and pestle and homogenized in 1.5 mL of extraction buffer containing 0.1 M phosphate buffer (pH 7.5) with 0.5 mM EDTA. The homogenate was centrifuged at 15,000×g for 30 minutes at 4°C, and the resulting clear supernatant was used to measure enzyme activity following standard protocols (Senthilkumar *et al*., 2021). Proline content was measured using 100 mg of ground plant tissue (shoot and root separately) according to Bates *et al*., (1973).

### Quantification of ethylene emission

To quantify ethylene, seedlings were grown in 20 mL glass vials (closed with rubber septum) with four seedlings per vial. Five-day-old seedlings were subjected to heat stress by exposing to 45°C for 4 hours, while seedlings in untreated vials served as controls. Immediately after heat treatment, a 1 mL aliquot of headspace gas was collected from each vial using a syringe through the rubber septum. Ethylene concentration was determined by injecting the collected gas into a gas chromatograph (GC-17A, Shimadzu) equipped with a Porapak T column and a flame ionization detector.

### Quantification of jasmonic acid

To measure jasmonic acid levels, five-days-old seedlings were exposed to heat stress at 45°C for a duration of four hours. Right after the heat exposure, the shoot tissues were collected using liquid nitrogen and then stored at -80°C until further use. The frozen samples were kept for lyophilization for three days using an Alpha 3-4 LSCbasic (CHRIST) lyophilizer. For hormone analysis, 25 mg of finely ground and weighed lyophilized leaf material was extracted with 1.5 mL of methanol, which included 40 ng of d6-JA as internal standards (Vadassery et al., 2012). The quantification of JA was performed using an Exion LC system connected to a Triple Quad 6500+ (Sciex). Samples were loaded onto a Zorbax Eclipse XDB-C18 (50x4.6mm, 1.8um, Agilent Technologies) column with a flow rate of 1.1 ml/min. The mobile phases A and B consisted of 0.05% formic acid in water and acetonitrile, respectively. The elution profile was set as follows: 0–0.5 min, 10% B; 0.5–4 min, 90% B; 4-4.02 min, 100% B; 4.02-4.5 min, 100% B; 4.5-4.51 min, 10% B; and from 4.51-7 min, 10% B. The column temperature was kept at 27°C. The Triple Quad 6500+ operated in negative ionization mode with an Ion Spray Voltage of -4500 eV, CUR gas at 45 psi, CAD set to medium, Temperature at 650, and GS1 and GS2 at 60 psi. Scheduled Multiple-reaction monitoring (MRM) was employed to track the analyte parent ion → product ion with a detection window of 60 seconds. Analyst 1.7 software (Applied Biosystems) facilitated data acquisition and processing. The phytohormone JA was quantified in relation to the signal of the corresponding internal standard.

### Transcriptome analysis

For RNA sequencing, five-day-old plants were harvested in liquid nitrogen immediately post-exposure to heat stress of 45°C for 4 h. Seedlings not exposed to the heat treatment were harvested at the same time and served as control. Shoots and roots were quickly separated and flash-frozen in liquid nitrogen. Samples were ground using mortar and pestle and 100 mg powder was collected in 2 ml Eppendorf safe lock tubes. Total RNA was extracted from plant samples using the PureLink™ RNA Mini Kit (Invitrogen) following the manufacturer’s protocol. The RNA integrity number (RIN) and concentration were checked using an Agilent 2100 Bioanalyzer (Agilent Technologies, Inc., Santa Clara, CA, USA). About 2 μg of total RNA was used for cDNA library preparation and sequencing. The paired-end sequencing libraries were prepared using Illumina HiSeq2000 RNA Library Preparation Kit as per manufacturer’s protocol (Illumina®, San Diego, CA). Quality control (QC) was done using FASTQC software. Adapter trimming was done using Trim Galore software, which removes the adapter sequences specific to Illumina sequencing and removes sequences that became shorter than 20 bp from a FASTQ file. Post-trimming, QC was performed again for all the samples. From the publicly available rice genome database Rice Genome Hub, the rice reference genome (*Oryza*_*sativa*_aus_N22.assembly) and gene model annotation files were downloaded. HISAT2 was used for indexing of the reference genome and alignment of the paired-end clean reads. HTSeq was used to retrieve the read counts mapped to each rice gene. Differential gene expression analysis between control and heat-stressed seedlings was carried out using the DESeq2 R package. Genes with adjusted P-value less than 0.05 and cut-off of Log_2_FC ≥ 1.5 were used for selecting upregulated genes, and Log_2_FC ≤ −1.5 was used for selecting downregulated genes.

### Quantitative real-time PCR analysis

RNA (2μg) was treated with DNase (Turbo DNase Kit) to eliminate genomic DNA contamination and quantified using NanoDrop™ One/OneC Microvolume UV-Vis Spectrophotometer (Thermo Scientific). First-strand cDNA synthesis was carried out using the Thermo Scientific RevertAid H Minus First Strand cDNA Synthesis Kit. Gene-specific primers for quantitative real-time PCR (qRT-PCR) were designed using QUANTPRIME, a primer design tool for high-throughput qPCR (Arvidsson *et al*., 2008). Expression analysis of all the candidate genes was performed using the SYBR Green assay (POWER SYBR® GREEN PCR Master Mix) with forward and reverse primers and cDNA templates on a BIO-RAD CFX384 and CFX96 Real-Time PCR System. Melt curve analysis was conducted to confirm primer specificity and binding efficiency. The housekeeping gene *OsACTIN11* was used as an internal control to normalise target gene expression across samples. Relative gene expression was calculated by converting Ct values into Log_2_ fold change (log_2_FC) values. All the primers used are listed in the supplementary table (**Table S1.5)**.

### Gene enrichment analysis

The Rice Genome Hub database was used for converting N22 gene IDs (A2J10) to Os IDs. Gene Ontology (GO) enrichment analysis for the identified differentially expressed genes (DEGs) was performed using ShinyGO (Ver 0.76) with an FDR cutoff of 0.05. The data obtained were plotted using ggplot2 R package. The GO child terms were manually classified under distinct broader GO terms.

### TF family enrichment and TF-Target identification

From the total DEGs, transcription factors (TFs) were identified by using all the nucleotide sequences as input into the TF database iTAK. For checking percentage enrichment, the number of DEG TFs (DETFs) in each family was divided by the total TFs in each family in rice. To identify the possible TF-target relations, we used the Regulation prediction option in the PlantRegMap database (Tian *et al*., 2019). It scans the 1kb promoter (upstream) sequence that we retrieved from RAP-DB database (Sakai et al., 2013), and infers regulatory interactions between TFs and DEGs and finds over-represented upstream TFs for the DEGs. All the DETFs that have enriched binding sites in the 1kb promoter region of DEGs were selected. We then estimated the Pearson correlation coefficient for the total TF-Target pairs based on their expression profiles (FPKM values) all samples and considered TF-target pairs with a FDR<0.05 for further analysis.

### TF-Target network analysis and module detection

The TF-target pairs thus identified in shoot and root were used as input to reconstruct, visualize and analyze the gene regulatory networks using Cytoscape 3.9.1. The TF-target pairs in both shoot and root were clustered into modules based on a topological clustering algorithm ‘GLay community clustering algorithm’ available in Cytoscape plugin clusterMaker2 (Shannon *et al*., 2003; Morris *et al*., 2011). Gene Ontology enrichment analysis was performed for the set of genes in each module using the GO-slim tool in the Panther database (Thomas *et al*., 2022). The number of TFs and percentage of heat stress-responsive genes in each module were identified by using the STIFDB database (v.2). To identify transcription factor-target gene (TF-Target) interactions as intramodular or intermodular, we compared the module assignments of each TF and its corresponding Target. If both belonged to the same module, the interaction was classified as intramodular; otherwise, it was considered intermodular.

### Identification of core hubs in functional interaction network

Cytoscape and the plugins STRING and clusterMaker2 were employed to construct the core hub network for the functional interactions. Functional interactions pertaining to all shoot and root DEGs were extracted from the STRING data network, and the total functional network was examined by the Analyze Network function to obtain various network metrices. K-means clustering was performed using Euclidean distance as the similarity metric and employing node attributes including betweenness centrality, closeness centrality, average shortest path length, clustering coefficient, degree and eccentricity. The analysis was conducted over 300 iterations, following the core-hub identification approach described by Kundariya *et al*., (2022). Genes within the cluster exhibiting the highest overall centrality, as measured by betweenness centrality and closeness centrality, as well as degree, were identified as core hubs. These genes were subsequently subjected to functional enrichment analysis utilizing the string enrichment function, with a false discovery rate (FDR) threshold of less than 0.05.

### Statistical analysis

Phenotyping, sectioning, staining experiments were all performed in more than three independent biological replicates times unless it’s mentioned otherwise. For all qRT-PCR experiments, a pairwise Student’s t-test was used to compare the gene expression of the heat-stressed and control groups. The student’s t-test was performed using R. Differences in continuous and discrete distributions were assessed using the non-parametric Wilcoxon rank sum test and Fisher’s Exact test respectively. Correction for multiple testing was done using False Discovery Rate (FDR). Enrichment of common DEGs between shoot and root was assessed using a permutation test. In each permutation, every common DEG was replaced with a random gene from the total expressed genes. The number of such randomly obtained genes that overlapped with the common DEG list was noted for 10,000 randomizations. From this, we estimated the Z-score, which indicates the distance of the actual observation to the mean of random expectation in terms of the number of standard deviations. P-values were estimated as the ratio of the randomly observed common genes greater than or equal to the number of observed common DEGs to the total number of random samples (10,000). All statistical analyses were done using R.

## Acknowledgements

ADA thanks the USDA-FAS Norman E Borlaug award for the financial support to conduct the RNA sequencing experiments. We thank IISER Tirupati core funding (ADA, AKS, R.V.K., and S.C.) and Ramalingaswami Re-entry Fellowship BT/RLF/Re-entry/05/2018 from Department of Biotechnology, Government of India (R.V.K., and S.C.). We acknowledge Prof. Y. Sreelakshmi and the RTGR facility funded by DBT SAHAJ (BT/INF/22/SP44787/2021) for the GC/MS analysis. We acknowledge Dr. V. Jyothilakshmi and the NIPGR Metabolome facility funded by DBT (BT/ INF/22/SP28268/2018) for LC-MS/MS analysis. We also acknowledge Darshan Damodaran and Devidutta Samantaray for their valuable assistance with laboratory experiments/discussions.

## Author contributions

ADA conceived the idea and supervised the work. AKS performed the phenotyping experiments and data analysis. SV performed the phenotyping experiments, staining, sectioning and biochemical analysis. TG performed the ACC/MeJA treatment experiments and qPCR analysis. AKS and ADA wrote the initial manuscript draft. ADA, SC and AKS were involved in designing the computation analysis, generation of figures, interpreting the data and editing the manuscript. RVK was involved in data analysis and editing of the manuscript. FBF was involved in editing the manuscript.

## Data and code availability

The raw transcriptome data will be submitted to the Sequence Read Archive (SRA) of the National Centre for Biotechnology Information (NCBI).

---

**Figure S1.**
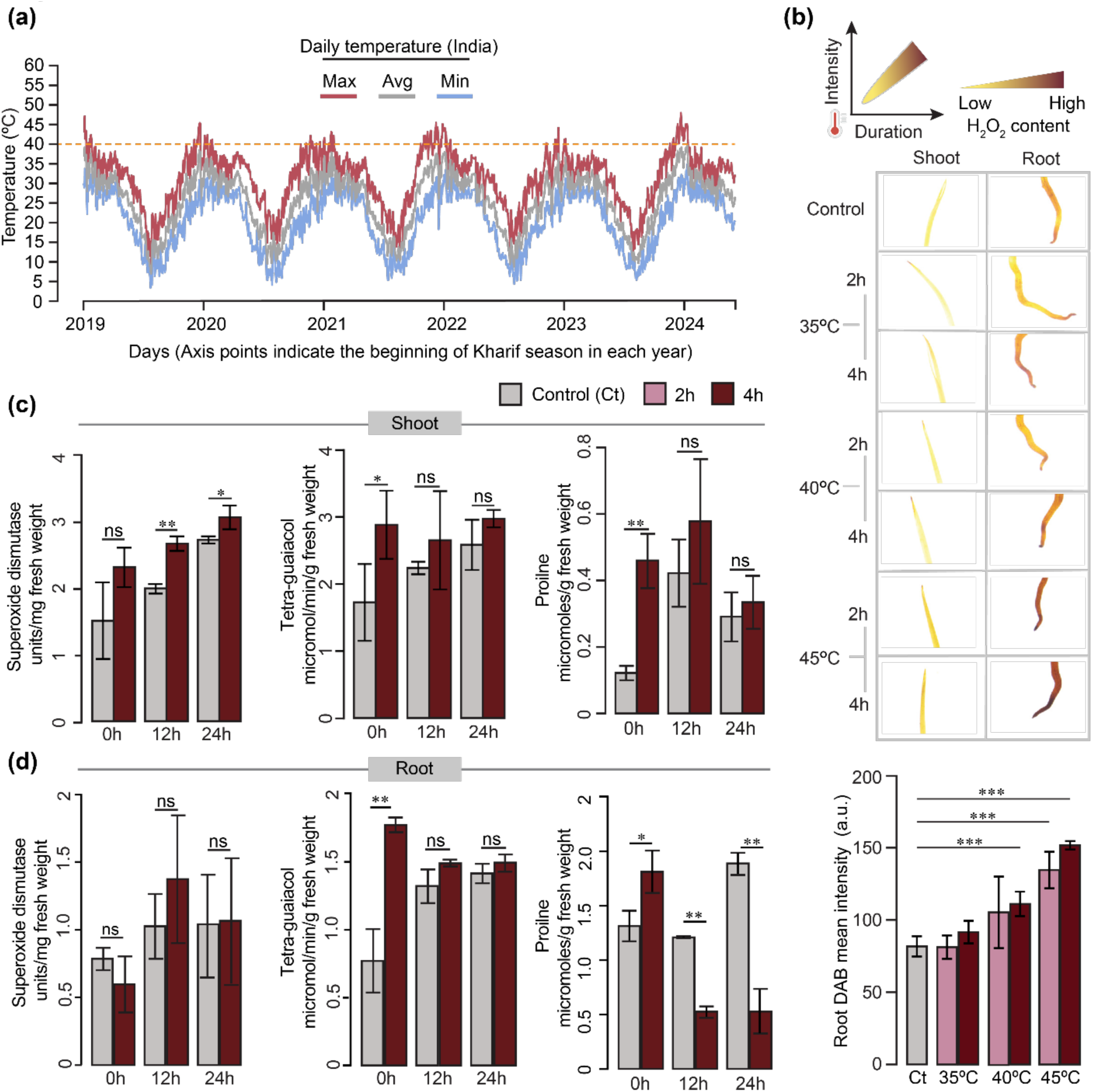
Biochemical responses of shoot and root under heat stress. **(a)** Daily temperatures (minimum, average and maximum) recorded in India in the past 6 years. Beginning of Kharif crop season-June is marked as a reference for each year on the Y axis. **(b)** H₂O₂ accumulation in shoot and root tissues was visualised using DAB staining immediately after heat stress. The intensity of colour represents H₂O₂ content, with gradients from low (yellow) to high (brown) levels. Rows show H₂O₂ accumulation across different temperatures and exposure durations. *n* = 8 for DAB quantification (mean ± SD is shown). **(c-d)** Antioxidant enzyme activity in shoots and roots: Superoxide dismutase (SOD) and Guaiacol peroxidase (GPOX) activities were measured at different time points post heat stress exposure. Proline content in shoot and root measured at 0, 12, and 24 hours (h) under control and heat stress conditions. *n*= 15 seedlings per biological replicate, with 3 independent biological replicates. Statistical analyses were performed using Student’s t-test, with specific P values shown for significant comparisons. **P < 0.05; ***P < 0.005; ns, not significant.

**Figure S2.**
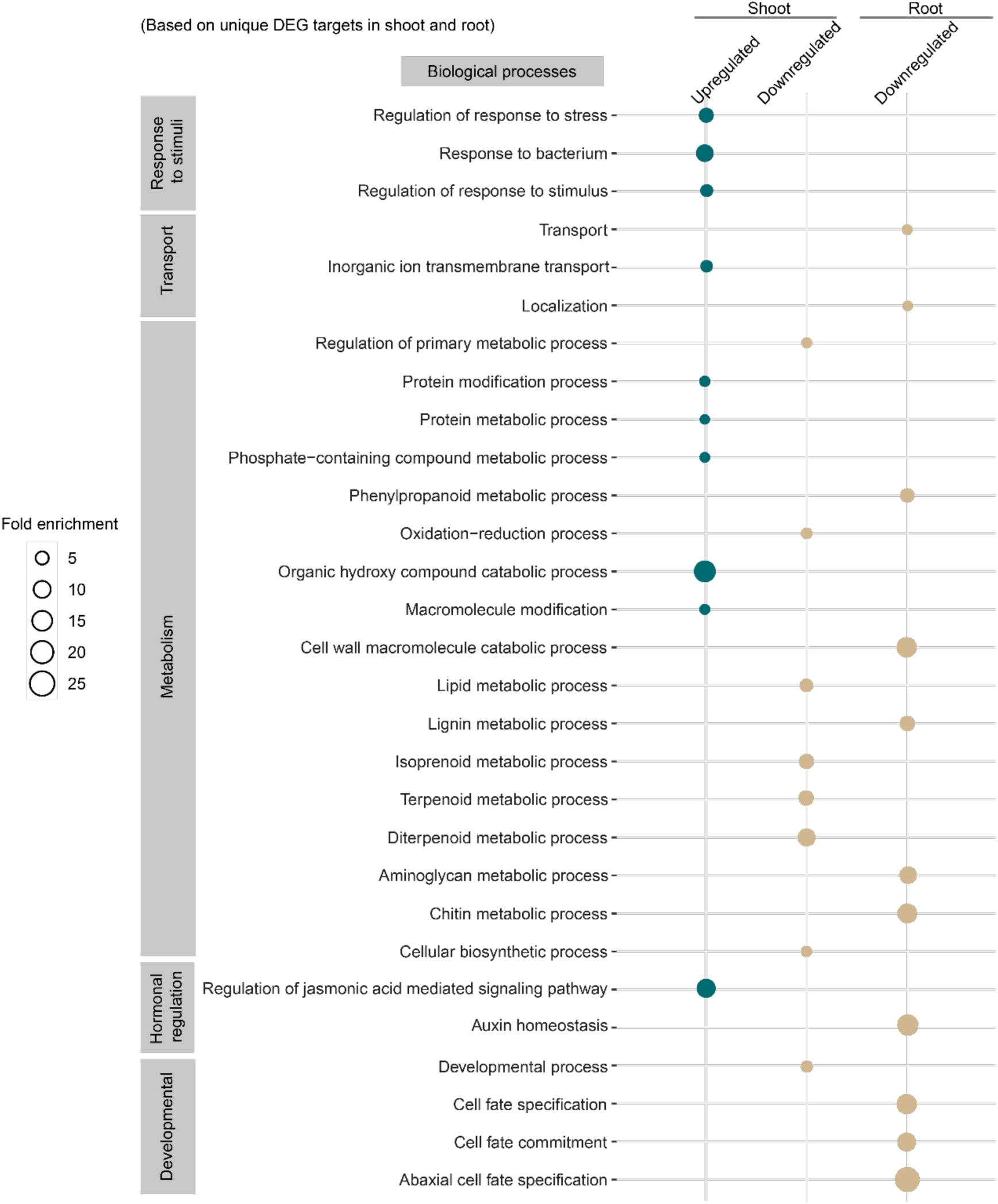
Biological processes enriched for differentially expressed genes (DEGs) unique in shoot and root tissues. The dot plot displays biological processes enriched for unique DEGs in shoot and root in response to heat stress. Processes are categorised into upregulated and downregulated groups for both shoot and root. The statistical significance of enrichment was determined using FDR-corrected p-values, with fold enrichment represented by dot size and significance level by colour intensity.

**Figure S3.**
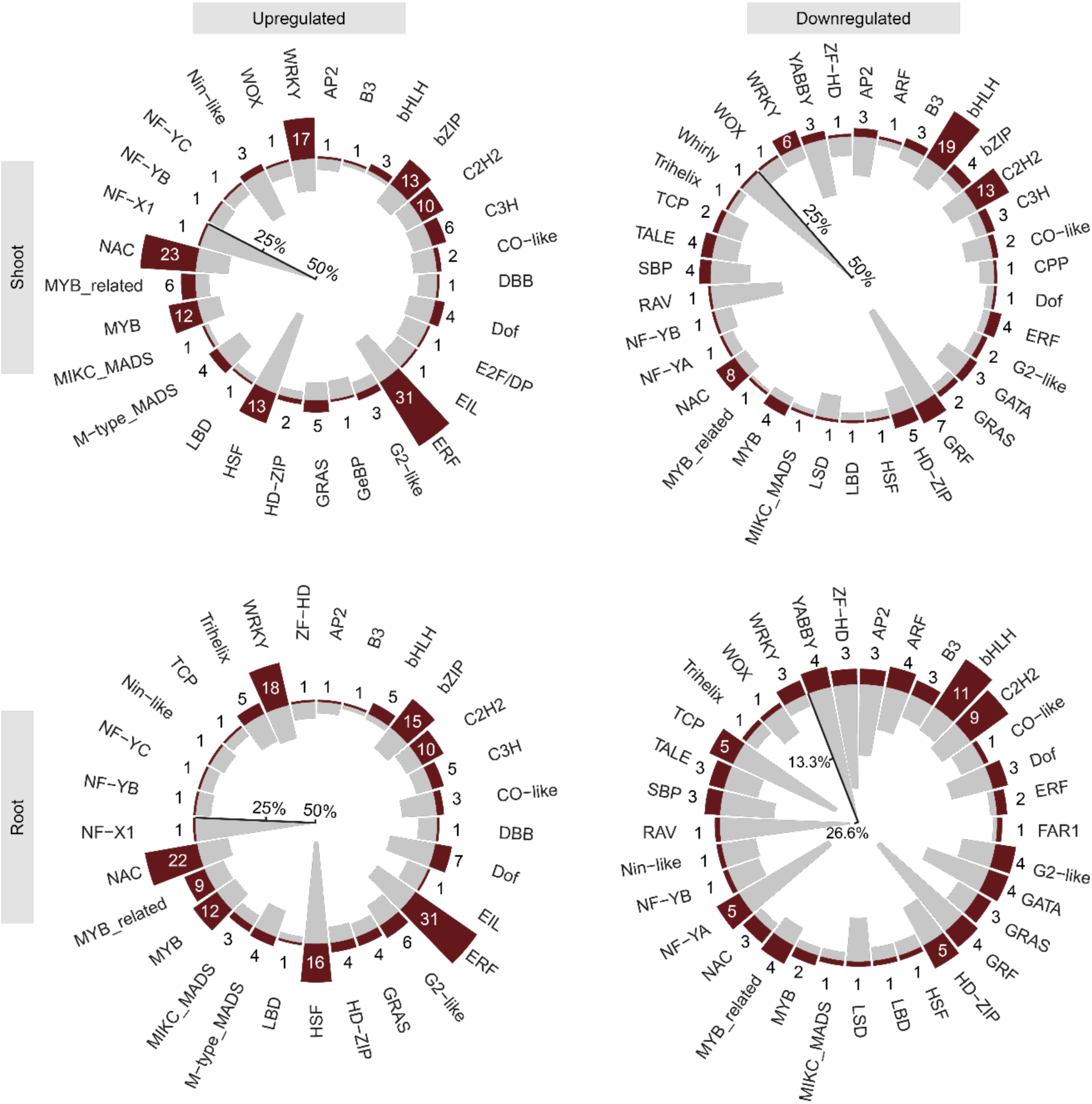
Distribution of transcription factor (TF) families in upregulated and downregulated DEGs in shoot and root. Circular plot showing the distribution of TF families among upregulated and downregulated DEGs in the shoot and root. The number of TFs within each family in *Oryza sativa* that are differentially expressed is indicated by the outer bars, and the inner bars indicate the percentage of gene expression from each TF family.

**Figure S4.**
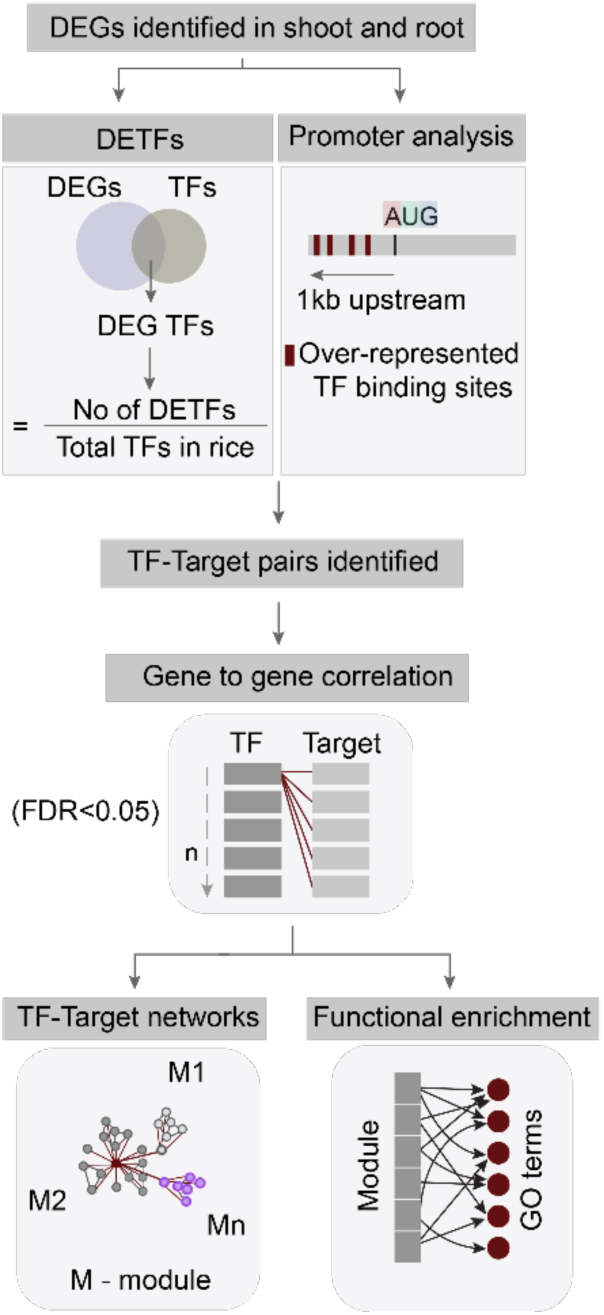
Workflow illustrating the TF-target networks and their functional enrichment. Acute heat stress responsive TF-Target network building includes DETF enrichment, promoter analysis, TF-target pair identification, gene-to-gene correlation analysis, network generation, module identification and functional enrichment

**Figure S5.**
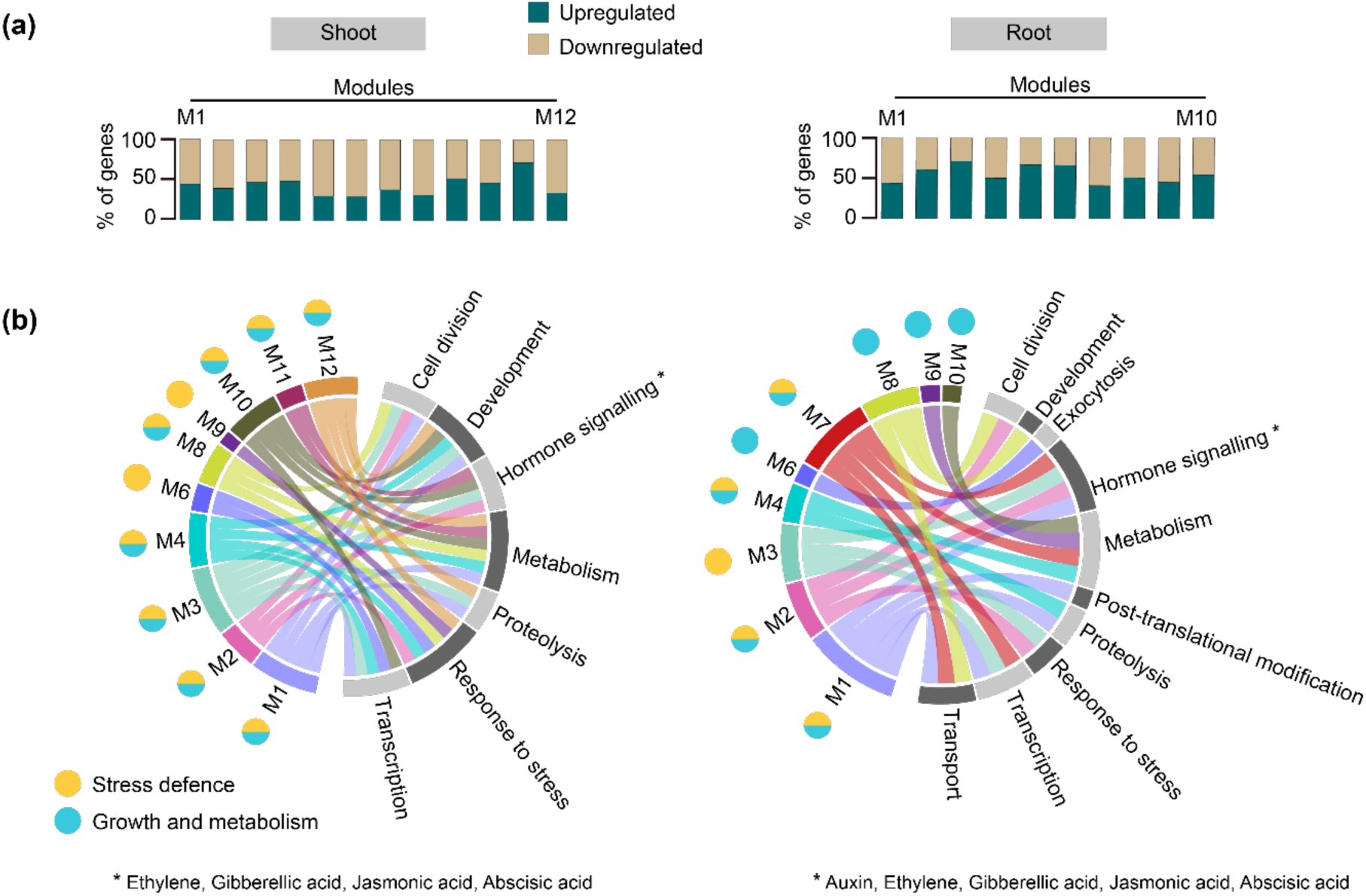
TF-Target modules regulate diverse biological processes in shoot and root. **(a)** Percentage of up(down)regulated genes in each module in shoot and root. **(b)** Circhos plot showing the association of TF modules with the biological processes in shoot and root. Biological processes are broadly categorized into two parent categories - stress defence and growth and metabolism; modules regulating these processes are indicated by respective coloured (yellow and blue) circles adjacently.

**Figure S6.**
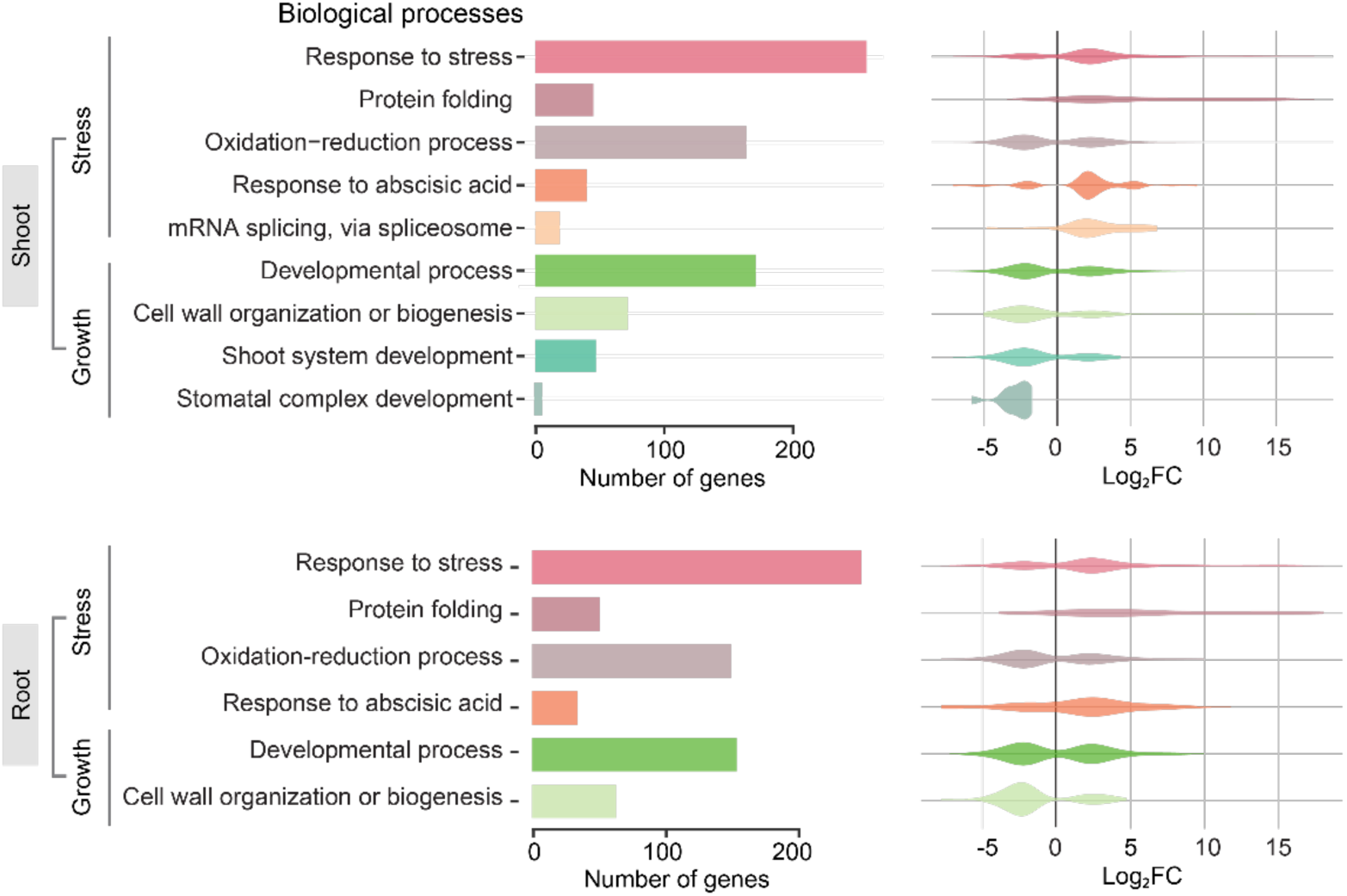
Functional enrichment of the DEGs functional network. Biological processes (BP) enriched in the total functional interactome network derived from the DEGs in shoot and root. The bar plot displays the number of genes associated with each biological process. Violin plots on the right show the distribution of log_2_ fold changes (FC) for genes involved in each of the processes.

## References

Albertos P, Dündar G, Schenk P, Carrera S, Cavelius P, Sieberer T, Poppenberger B. 2022. Transcription factor BES1 interacts with HSFA1 to promote heat stress resistance of plants. The EMBO Journal 41: e108664.

Alshareef NO, Otterbach SL, Allu AD, Woo YH, de Werk T, Kamranfar I, Mueller-Roeber B, Tester M, Balazadeh S, Schmöckel SM. 2022. NAC transcription factors ATAF1 and ANAC055 affect the heat stress response in Arabidopsis. Scientific Reports 12: 11264.

Arvidsson S, Kwasniewski M, Riaño-Pachón DM, Mueller-Roeber B. 2008. QuantPrime – a flexible tool for reliable high-throughput primer design for quantitative PCR. BMC Bioinformatics 9: 465.

Aryan S, Gulab G, Habibi N, Kakar K, Sadat MI, Zahid T, Rashid RA. 2022. Phenological and physiological responses of hybrid rice under different high-temperature at seedling stage. Bulletin of the National Research Centre 46: 45.

Bakery A, Vraggalas S, Shalha B, Chauhan H, Benhamed M, Fragkostefanakis S. 2024. Heat stress transcription factors as the central molecular rheostat to optimize plant survival and recovery from heat stress. New Phytologist 244: 51–64.

Begcy, K., Sandhu, J. and Walia, H., 2018. Transient heat stress during early seed development primes germination and seedling establishment in rice. Frontiers in Plant Science, 9, p.1768.

Cai H, Wang H, Zhou L, Li B, Zhang S, He Y, Guo Y, You A, Jiao C, Xu Y. 2023. Time-series transcriptomic analysis of contrasting rice materials under heat stress reveals a faster response in the tolerant cultivar. International Journal of Molecular Sciences 24: 9408.

Calvin K, Dasgupta D, Krinner G, Mukherji A, Thorne PW, Trisos C, Romero J, Aldunce P, Barrett K, Blanco G, et al. 2023. IPCC, 2023: Climate Change 2023: Synthesis Report. Contribution of Working Groups I, II and III to the Sixth Assessment Report of the Intergovernmental Panel on Climate Change [Core Writing Team, H. Lee and J. Romero (eds.)]. IPCC, Geneva, Switzerland.

Casal JJ, Balasubramanian S. 2019. Thermomorphogenesis. Annual Review of Plant Biology 70: 321–346.

Chao L-M, Liu Y-Q, Chen D-Y, Xue X-Y, Mao Y-B, Chen X-Y. 2017. Arabidopsis transcription factors SPL1 and SPL12 confer plant thermotolerance at reproductive stage. Molecular Plant 10: 735–748.

Dey S, Corina Vlot A. 2015. Ethylene responsive factors in the orchestration of stress responses in monocotyledonous plants. Frontiers in Plant Science 6.

Facelli JM. 2008. Specialized seedling strategies I: seedlings in stressful environments. In: Seedling Ecology and Evolution. Cambridge University Press, 56–78.

Freyre-González JA, Manjarrez-Casas AM, Merino E, Martinez-Nuñez M, Perez-Rueda E, Gutiérrez-Ríos R-M. 2013. Lessons from the modular organization of the transcriptional regulatory network of *Bacillus subtilis*. BMC Systems Biology 7: 127.

Gauthey A, Kahmen A, Limousin J, Vilagrosa A, Didion-Gency M, Mas E, Milano A, Tunas A, Grossiord C. 2024. High heat tolerance, evaporative cooling, and stomatal decoupling regulate canopy temperature and their safety margins in three European oak species. Global Change Biology 30: e17439.

González-Schain N, Dreni L, Lawas LMF, Galbiati M, Colombo L, Heuer S, Jagadish KSV, Kater MM. 2016. Genome-wide transcriptome analysis during anthesis reveals new insights into the molecular basis of heat stress responses in tolerant and sensitive rice varieties. Plant and Cell Physiology 57: 57–68.

Guo M, Liu J-H, Ma X, Luo D-X, Gong Z-H, Lu M-H. 2016. The plant heat stress transcription factors (HSFs): structure, regulation, and function in response to abiotic stresses. Frontiers in Plant Science 7.

Han F, Chen H, Li X-J, Yang M-F, Liu G-S, Shen S-H. 2009. A comparative proteomic analysis of rice seedlings under various high-temperature stresses. Biochimica et Biophysica Acta (BBA) - Proteins and Proteomics 1794: 1625–1634.

Hasanuzzaman M, Nahar K, Alam Md, Roychowdhury R, Fujita M. 2013. Physiological, biochemical, and molecular mechanisms of heat stress tolerance in plants. International Journal of Molecular Sciences 14: 9643–9684.

He Y, Guan H, Li B, Zhang S, Xu Y, Yao Y, Yang X, Zha Z, Guo Y, Jiao C, et al. 2023. Transcriptome analysis reveals the dynamic and rapid transcriptional reprogramming involved in heat stress and identification of heat response genes in rice. International Journal of Molecular Sciences 24(19): 14802.

He Z, Webster S, He SY. 2022. Growth–defense trade-offs in plants. Current Biology 32: R634–R639.

Hoagland DR, Arnon DI. 1950. The water-culture method for growing plants without soil.

Hoang XLT, Nhi DNH, Thu NBA, Thao NP, Tran L-SP. 2017. Transcription factors and their roles in signal transduction in plants under abiotic stresses. Current genomics 18: 483–497.

Huang H, Ullah F, Zhou D-X, Yi M, Zhao Y. 2019. Mechanisms of ROS regulation of plant development and stress responses. Frontiers in Plant Science 10.

Huang J, Zhang F, Xue Y, Lin J. 2017. Recent changes of rice heat stress in Jiangxi province, southeast China. International Journal of Biometeorology 61: 623–633.

Huang J, Zhao X, Bürger M, Chory J, Wang X. 2023. The role of ethylene in plant temperature stress response. Trends in Plant Science 28: 808–824.

Huot B, Yao J, Montgomery BL, He SY. 2014. Growth–defense tradeoffs in plants: a balancing act to optimize fitness. Molecular Plant 7: 1267–1287.

Iqbal N, Fatma M, Khan NA, Umar S. 2019. Regulatory role of proline in heat stress tolerance. In: Plant Signaling Molecules. Elsevier, 437–448.

Jagadish SVK, Muthurajan R, Oane R, Wheeler TR, Heuer S, Bennett J, Craufurd PQ. 2010. Physiological and proteomic approaches to address heat tolerance during anthesis in rice (*Oryza sativa* L.). Journal of Experimental Botany 61: 143–156.

Janni M, Gullì M, Maestri E, Marmiroli M, Valliyodan B, Nguyen HT, Marmiroli N. 2020. Molecular and genetic bases of heat stress responses in crop plants and breeding for increased resilience and productivity. Journal of Experimental Botany 71: 3780–3802.

Jung K-H, Ko H-J, Nguyen MX, Kim S-R, Ronald P, An G. 2012. Genome-wide identification and analysis of early heat stress responsive genes in rice. Journal of Plant Biology 55: 458– 468.

Kacprzyk J, McCabe PF, Ng CK -Y. 2025. Beat the heat: need for research studying plant cell death induced by extreme temperatures. New Phytologist. 10.1111/nph.70045.

Kim M-S, Kim D, Kang NS, Kim J-R. 2017. The core regulatory network in human cells. Biochemical and Biophysical Research Communications 484: 348–353.

Kundariya H, Sanchez R, Yang X, Hafner A, Mackenzie SA. 2022. Methylome decoding of RdDM-mediated reprogramming effects in the Arabidopsis MSH1 system. Genome Biology 23: 167.

Licausi, F., Ohme-Takagi, M. and Perata, P., 2013. APETALA 2/Ethylene Responsive Factor (AP2/ERF) transcription factors: Mediators of stress responses and developmental programs. New Phytologist 199(3): 639–649.

Li H, Liu Y, Li Y, Yang Q, Yang T, Zhou Z, Li Y, Zhang N, Lyu Y, Zhu Y, et al. 2023.Heat shock transcription factors regulate thermotolerance gene networks in tomato (*Solanum lycopersicum*) flower buds. Horticultural Plant Journal 11(1): 199–210.

Liu Y, Liu X, Wang X, Gao K, Qi W, Ren H, Hu H, Sun D, Bai J, Zheng S. 2020. Heterologous expression of heat stress-responsive AtPLC9 confers heat tolerance in transgenic rice. BMC Plant Biology 20: 514.

Liu S, Zeng F, Fan G, Dong Q. 2021. Identification of hub genes and construction of a transcriptional regulatory network associated with tumor recurrence in colorectal cancer by weighted gene co-expression network analysis. Frontiers in Genetics 12.

Mangrauthia SK, Bhogireddy S, Agarwal S, Prasanth V V., Voleti SR, Neelamraju S, Subrahmanyam D. 2017. Genome-wide changes in microRNA expression during short and prolonged heat stress and recovery in contrasting rice cultivars. Journal of Experimental Botany 68: 2399–2412.

Mittal D, Madhyastha DA, Grover A. 2012. Genome-wide transcriptional profiles during temperature and oxidative stress reveal coordinated expression patterns and overlapping regulons in rice. PLoS ONE 7: e40899.

Morris JH, Apeltsin L, Newman AM, Baumbach J, Wittkop T, Su G, Bader GD, Ferrin TE. 2011. clusterMaker: a multi-algorithm clustering plugin for Cytoscape. BMC Bioinformatics 12: 436.

Müller-Dott S, Tsirvouli E, Vazquez M, Ramirez Flores RO, Badia-i-Mompel P, Fallegger R, Türei D, Lægreid A, Saez-Rodriguez J. 2023. Expanding the coverage of regulons from high-confidence prior knowledge for accurate estimation of transcription factor activities. Nucleic Acids Research 51: 10934–10949.

Müller M, Munné-Bosch S. 2015. Ethylene Response Factors: A key regulatory hub in hormone and stress signaling. Plant Physiology 169: 32–41.

Naika M, Shameer K, Mathew OK, Gowda R, Sowdhamini R. 2013. STIFDB2: An updated version of plant stress-responsive transcription factor data base with additional stress signals, stress-responsive transcription factor binding sites and stress-responsive genes in arabidopsis and rice. Plant and Cell Physiology 54(2):e8.

Ohama N, Sato H, Shinozaki K, Yamaguchi-Shinozaki K. 2017. Transcriptional regulatory network of plant heat stress response. Trends in Plant Science 22: 53–65.

Ravindra, K., Bhardwaj, S., Ram, C., Goyal, A., Singh, V., Venkataraman, C., Bhan, S.C., Sokhi, R.S. and Mor, S., 2024. Temperature projections and heatwave attribution scenarios over India: A systematic review. Heliyon 10(4).

Reynoso MA, Borowsky AT, Pauluzzi GC, Yeung E, Zhang J, Formentin E, Velasco J, Cabanlit S, Duvenjian C, Prior MJ, et al. 2022. Gene regulatory networks shape developmental plasticity of root cell types under water extremes in rice. Developmental Cell 57(9): 1177–1192.e6.

Sakai H, Lee SS, Tanaka T, Numa H, Kim J, Kawahara Y, Wakimoto H, Yang C, Iwamoto M, Abe T, et al. 2013. Rice Annotation Project Database (RAP-DB): An integrative and interactive database for rice genomics. Plant and Cell Physiology 54(2):e6.

Sarkar NK, Kim Y-K, Grover A. 2014. Coexpression network analysis associated with call of rice seedlings for encountering heat stress. Plant Molecular Biology 84: 125–143.

Senthilkumar M, Amaresan N, Sankaranarayanan A. 2021. Plant-Microbe Interactions. New York, NY: Springer US.

Shannon P, Markiel A, Ozier O, Baliga NS, Wang JT, Ramage D, Amin N, Schwikowski B, Ideker T. 2003. Cytoscape: A software environment for integrated models of biomolecular interaction networks. Genome Research 13: 2498–2504.

de Silva KK, Dunwell JM, Wickramasuriya AM. 2022. Weighted Gene Correlation Network Analysis (WGCNA) of Arabidopsis somatic embryogenesis (se) and identification of key gene modules to uncover se-associated hub genes. International Journal of Genomics 2022: 7471063.

Song L, Huang SC, Wise A, Castanon R, Nery JR, Chen H, Watanabe M, Thomas J, Bar-Joseph Z, Ecker JR. 2016. A transcription factor hierarchy defines an environmental stress response network. Science 354(6312): aag1550.

Sun Y, Oh D-H, Duan L, Ramachandran P, Ramirez A, Bartlett A, Tran K-N, Wang G, Dassanayake M, Dinneny JR. 2022. Divergence in the ABA gene regulatory network underlies differential growth control. Nature Plants 8: 549–560.

Surabhi G-K, Badajena B. 2020. Recent advances in plant heat stress transcription factors. In: Transcription factors for abiotic stress tolerance in plants. Elsevier, 153–200.

Suzuki N, Katano K. 2018. Coordination between ROS regulatory systems and other pathways under heat stress and pathogen attack. Frontiers in Plant Science 9.

Tan W, Chen J, Yue X, Chai S, Liu W, Li C, Yang F, Gao Y, Gutiérrez Rodríguez L, Resco de Dios V, et al. 2023. The heat response regulators HSFA1s promote *Arabidopsis* thermomorphogenesis via stabilizing PIF4 during the day. Science Advances 9(44): eadh1738.

Thomas, P.D., Ebert, D., Muruganujan, A., Mushayahama, T., Albou, L.P. and Mi, H., 2022. PANTHER: Making genome-scale phylogenetics accessible to all. Protein Science 31(1): 8–22.

Tian F, Yang D-C, Meng Y-Q, Jin J, Gao G. 2019. PlantRegMap: charting functional regulatory maps in plants. Nucleic Acids Research 48(D1):D1104–D1113.

Vadassery, J., Reichelt, M., Hause, B., Gershenzon, J., Boland, W. and Mithöfer, A., 2012. CML42-mediated calcium signaling coordinates responses to Spodoptera herbivory and abiotic stresses in Arabidopsis. Plant physiology 159(3): 1159–1175.

Vecellio DJ, Kong Q, Kenney WL, Huber M. 2023. Greatly enhanced risk to humans as a consequence of empirically determined lower moist heat stress tolerance. Proceedings of the National Academy of Sciences 120(42): e2305427120.

Wang FZ, Niyogi KK. 2025. Towards targeted engineering of promoters via deletion of repressive *cis*-regulatory elements. New Phytologist 245: 1805–1807.

Wang X, Tan NWK, Chung FY, Yamaguchi N, Gan E-S, Ito T. 2023. Transcriptional regulators of plant adaptation to heat stress. International Journal of Molecular Sciences 24: 13297.

Wang Y, Zhang Y, Zhang Q, Cui Y, Xiang J, Chen H, Hu G, Chen Y, Wang X, Zhu D, et al. 2019. Comparative transcriptome analysis of panicle development under heat stress in two rice (*Oryza sativa* L.) cultivars differing in heat tolerance. PeerJ 7: e7595.

Wassmann R, Jagadish SVK, Sumfleth K, Pathak H, Howell G, Ismail A, Serraj R, Redona E, Singh RK, Heuer S. 2009. Chapter 3: Regional vulnerability of climate change impacts on asian rice production and scope for adaptation. In: 91–133.

Wilkins O, Hafemeister C, Plessis A, Holloway-Phillips M-M, Pham GM, Nicotra AB, Gregorio GB, Jagadish SVK, Septiningsih EM, Bonneau R, et al. 2016. EGRINs (Environmental Gene Regulatory Influence Networks) in rice that function in the response to water deficit, high temperature, and agricultural environments. The Plant Cell 28: 2365–2384.

WMO. 2024. July sets new temperature records.

Wu T-Y, Goh H, Azodi CB, Krishnamoorthi S, Liu M-J, Urano D. 2021. Evolutionarily conserved hierarchical gene regulatory networks for plant salt stress response. Nature Plants 7: 787–799.

Wu J, Liu P, Liu Y. 2023. Thermosensing and thermal responses in plants. Trends in Biochemical Sciences 48: 923–926.

Xiang Q, Rathinasabapathi B. 2022. Differential tolerance to heat stress of young leaves compared to mature leaves of whole plants relate to differential transcriptomes involved in metabolic adaptations to stress. AoB PLANTS 14(4)plac024.

Yang J, Chen X, Zhu C, Peng X, He X, Fu J, Ouyang L, Bian J, Hu L, Sun X, et al. 2015. Using rna-seq to profile gene expression of spikelet development in response to temperature and nitrogen during meiosis in rice (*Oryza sativa* L.). PLOS ONE 10: e0145532.

Yang Y, Zhang C, Zhu D, He H, Wei Z, Yuan Q, Li X, Gao X, Zhang B, Gao H, et al. 2022. Identifying candidate genes and patterns of heat-stress response in rice using a genome-wide association study and transcriptome analyses. The Crop Journal 10: 1633–1643.

Yin L, Zander M, Huang SC, Xie M, Song L, Guzmán JPS, Hann E, Shanbhag BK, Ng S, Jain S, et al. 2023. Transcription factor dynamics in cross-regulation of plant hormone signaling pathways. BioRxiv. doi: 10.1101/2023.03.07.531630.

Zhang H, Zhou J-F, Kan Y, Shan J-X, Ye W-W, Dong N-Q, Guo T, Xiang Y-H, Yang Y-B, Li Y-C, et al. 2022. A genetic module at one locus in rice protects chloroplasts to enhance thermotolerance. Science 376: 1293–1300.

Zhou Z, Eichner C, Nilsen F, Jonassen I, Dondrup M. 2021. A novel approach to co-expression network analysis identifies modules and genes relevant for moulting and development in the Atlantic salmon louse (*Lepeophtheirus salmonis*). BMC Genomics 22(1):832.

